# Genomic classification and immune profiling define mucosal melanoma

**DOI:** 10.64898/2026.10.02.756347

**Authors:** Yingxiao Shi, Priyanka Sharma, Jia-ren Lin, Aikaterini Dedeilia, Tuulia Vallius, Giuseppe Tarantino, Roxanne Pelletier, Mariana L Lopez, Nil Aygün, Michael P Manos, Sonia Cohen, Aleigha R Lawless, Yi Daniel Lu, Madeline K Roelofs, Shishir M Pant, Jiajia Chen, David E Fisher, Rameen Beroukhim, Christine G Lian, Elizabeth I Buchbinder, Peter K Sorger, Megan L. Insco, David Liu, Genevieve Boland

## Abstract

Mucosal melanoma is a rare, lethal melanoma subtype with distinct genetics and limited benefit from immunotherapy. To define features associated with mucosal melanoma clinical outcomes, we assembled a large, multimodally profiled cohort spanning primary and metastatic disease. Compared with cutaneous melanoma, primary mucosal melanoma showed greater genomic heterogeneity, ploidy and aneuploidy, and a more dedifferentiated, proliferative transcriptional state. Analysis of genomic sequencing of 339 patients with mucosal melanoma identified four mutually exclusive genomic subtypes: CDK4/MDM2/CCND1, SF3B1-mutated, TP53/ATRX, and mucosal melanoma Triple WT, associated with distinct anatomic sites but similar outcomes. In the tumor microenvironment, we observed reduced T cell infiltration and increased CD163^+^ tumor-associated macrophages in mucosal melanoma relative to cutaneous melanoma. In matched longitudinal samples, CD163^+^ tumor-associated macrophages increased with disease progression and immunotherapy resistance. CD163^+^ tumor-associated macrophages engaged non-regulatory T cells through GAL9–TIM-3 interactions, nominating TIM-3 blockade as a therapeutic strategy for mucosal melanoma. These findings define the distinct aggressive biology of mucosal melanoma and highlight potential therapeutic targets and mechanisms of immune resistance.

## INTRODUCTION

Mucosal melanoma (MM) is a rare melanoma subtype arising from melanocytes within mucosal membranes^1^. Although derived from melanocytes, MM is biologically and clinically distinct from cutaneous melanoma (CM). Patients with MM have the lowest relative survival at three years of any melanoma subtype^2^. As MM most frequently arises in sinonasal, vulvovaginal, and anorectal tissues, which are more difficult to monitor than skin, it is not surprising that MM are diagnosed at more advanced stages^3^. MM is also more likely to become metastatic as compared to stage-matched CM^4^. An additional challenge in MM is its anatomic heterogeneity, which fragments clinical care across specialties, complicating tissue collection and standardization of guideline-based care. Finally, MM is substantially less responsive to treatment than CM across every therapeutic modality^5^. Current treatment strategies—including surgical resection^6^, targeted therapy^7–10^,and immunotherapy^11–14^—collectively yield substantially poorer outcomes compared with CM. Although immune checkpoint blockade (ICB) has transformed CM treatment, its efficacy in MM remains limited. In pooled nivolumab-treated patients, MM had lower ORRs than CM after PD-1 monotherapy (23% vs 41%) and PD-1/CTLA-4 blockade (37% vs 60%)^11,15^. Further, in MM, even when a response to PD-1 blockade is achieved, it is less durable than in CM^16^. Together, these findings underscore an urgent need to understand the mechanisms underlying MM metastatic potential and immunotherapy resistance to develop more effective treatment strategies.

Recent studies have begun to characterize the unique genomic landscape of MM, particularly in comparison with CM. Unlike CM, as MM develops in sun-protected sites, MM lacks ultraviolet (UV)-induced mutational signatures thus exhibiting a significantly lower tumor mutational burden. Instead, MM is characterized by extensive structural genomic instability, including frequent copy-number alterations and chromosomal rearrangements^17,18^. Lastly, MM harbors a distinct spectrum of genomic driver alterations^18–22^, including a lack of hotspot *BRAF* mutations (4.5-7% in MM vs. 40-50% in CM)^18,22^ and an enrichment of recurrent point mutations in *SF3B1* (∼35% in MM vs. ∼1% in CM)^19,23^. In addition, *CDK4* and *CCND1* are among the most amplified genes in MM and these alterations are enriched in MM (∼40%) vs. CM (∼9.6%)^20,24–26^, highlighting that cell cycle dysregulation may be important in MM pathogenesis^27^.

The gene expression landscape in MM has not yet been comprehensively characterized at scale, particularly across anatomic sites and between primary and metastatic tumors. While gene expression analyses in CM have demonstrated substantial tumor cell-state plasticity with dynamic transitions between melanocytic, neural crest-like, dedifferentiation, and mesenchymal states contributing to disease progression and resistance to ICB^28–31^, it remains unknown whether similar transcriptional programs define MM biology. Recent single-cell transcriptomic analysis from our group showed that MM tumor cells are enriched for migratory neural crest-associated programs as compared with CM, suggesting a more dedifferentiated and invasive phenotype of melanoma^22^. However, because these analyses were performed on a small number of predominantly post-ICB treated tumors, it remains unclear whether these tumor states represent intrinsic features of untreated MM or emerge as a shared feature of therapeutic resistance across melanoma subtypes.

Tumor-intrinsic alterations alone are unlikely to fully explain MM’s aggressive clinical behavior or poor response to immunotherapy. Increasing evidence suggests that the tumor microenvironment (TME) plays a critical role in disease progression and treatment response^32–35^. MM is thought to respond poorly to ICB in part because of its comparative "cold" immune microenvironment^36^. Prior studies have reported significantly reduced tumor-infiltrating lymphocytes (TILs) in MM relative to CM in primary tumors^37,38^, and single cell transcriptomic profiling of a limited number of patients showed increased infiltration of CXCL3^+^ tumor-associated macrophages (TAMs) in MM relative to acral melanoma (AM) and CM, suggesting that immunosuppressive macrophage populations may also contribute to immune evasion^39^. However, these studies were performed mainly on untreated primary or surgically resected tumors profiled at a single timepoint, and how the TME evolves during disease progression and immunotherapy resistance in MM is unknown.

To address these knowledge gaps, we assembled a large, multi-institutional cohort of patients with MM and performed an integrated multi-omic analysis incorporating whole-exome sequencing (WES), targeted sequencing, bulk RNA sequencing (RNA-seq), single-cell RNA sequencing (scRNA-seq), Orion multiplex imaging, and histopathological analysis. Using this integrated framework, we show that MM is intrinsically distinct from CM in primary tumors across anatomic sites; demonstrate for the first time a genomic classification of mutually exclusive mutational subtypes; and show that MM develops a macrophage-enriched immunosuppressive microenvironment that evolves during disease progression and immunotherapy resistance.

## RESULTS

### Primary MM is a unified cell-intrinsically distinct entity across anatomic sites

To systematically compare the intrinsic biology of MM and CM, we assembled a large multi-institutional cohort integrating genomic, transcriptomic, and spatial profiling. The cohort, experiment data generation, and analyses are summarized in **Fig. 1a** and **Extended Data Fig. 1a,b**, with sample-level clinical and molecular annotations are provided in **Supplementary Table 1**–**3**. Genomic profiling included targeted sequencing from Dana-Farber Cancer Institute (DFCI) (n = 86) and a multi-institutional cohort (AACR GENIE^40^) (n = 253) together with WES of 122 samples from 84 patients collected at Massachusetts General Hospital (MGH) and DFCI. Bulk RNA sequencing was performed on 72 tumors, overlapping with the WES cohort, and we leveraged a previously published scRNA-seq dataset from ten post-ICB-tumors generated as part of this collaboration^22^. Pathology-confirmed MM formalin-fixed, paraffin-embedded (FFPE) specimens were characterized using 16-plex Orion imaging^41^, enabling simultaneous detection of tumor cells (SOX10, MART1), proliferation (KI67), stromal structures (CD31, αSMA), and major immune populations (CD45, CD3E, CD4, CD8A, FOXP3, CD20, CD11c, and CD163) (Methods, **Extended Data Fig. 1c**).

**Fig. 1:**
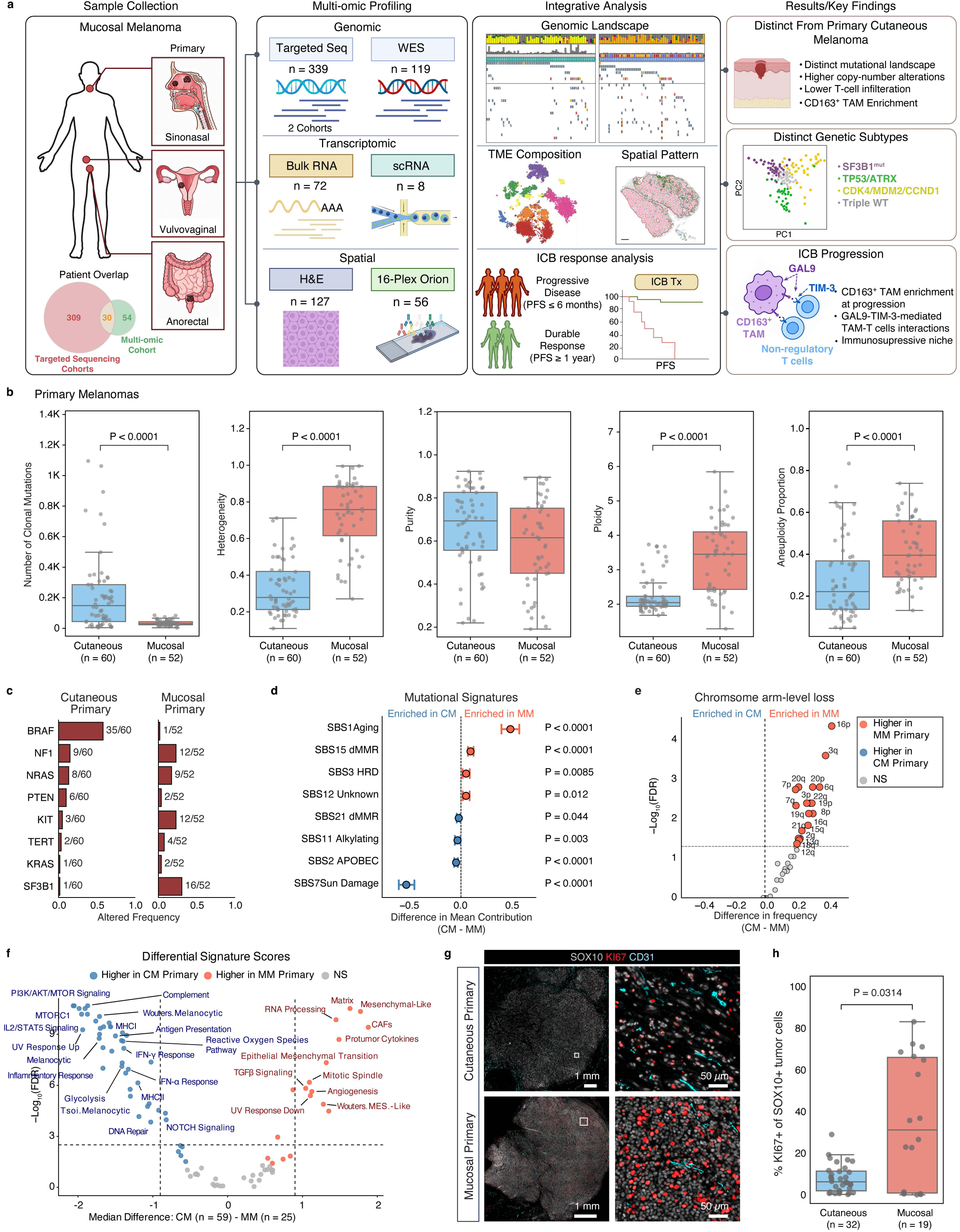
Study cohort overview and molecular characterization of MM. **a**, Overview of the patient cohort, summarizing primary tumor collection by anatomic site, the profiling strategies applied to each sample set and the key findings of the analysis. **b**, Genomic features comparison between primary CM (*n* = 60) and MM (*n* = 52): number of clonal mutations, intratumoral heterogeneity, purity, ploidy and aneuploidy. Box plots show the median and interquartile range (IQR), with whiskers extending to 1.5 ’IQR and individual tumors shown as points. Groups were compared using two-sided Mann-Whitney *U* tests; significant *P* values (*P* < 0.05) are shown. **c**, Recurrently altered driver genes in primary CM (TCGA) and primary MM. Significantly mutated genes were identified using MutSig2CV, and the frequency of genomic alterations is shown for each cohort. **d**, Comparison of SBS mutational signature activities between primary CM and primary MM. SBS signature contributions were inferred using deconstructSigs, and bars represent the mean contribution of each signature across samples. **e**, Comparison of chromosome arm-level copy-number loss between primary CM and MM. The *x* axis shows the difference in the proportion of tumors with arm-level loss (primary CM – primary MM) and the *y* axis shows the false discovery FDR-adjusted *P* value. *P* values were calculated using two-sided Fisher’s exact tests with Benjamini-Hochberg correction; the dashed line indicates an adjusted *P* value of 0.05. **f**, Differentially enriched transcriptomic signatures between primary CM and primary MM. Signature scores were calculated using Hallmark pathways, melanoma cell-state signatures, and published gene expression programs. The *x* axis shows the mean difference in signature score (primary CM − primary MM), and the *y* axis shows the FDR-adjusted *P* value. **g**, Representative Orion multiplex immunofluorescence images of primary CM and primary MM stained for SOX10 (white), KI67 (red), and CD31 (cyan). Right panels show higher-magnification views of the boxed regions. Scale bars, 1 mm and 50 μm. **h**, Quantification of proliferating SOX10⁺KI67⁺ tumor cells in primary CM (*n* = 32) and primary mucosal MM (*n* = 19). Box plots show the median and interquartile range (IQR), with whiskers extending to 1.5 ’IQR and individual tumors shown as points. Groups were compared using a two-sided Mann-Whitney *U* test.

To determine whether primary MM and CM are genetically distinct, we compared WES-derived genomic features from 52 primary MM tumors collected across different anatomic sites (**Extended Data Fig. 1d**) with primary CM tumors from TCGA^42^. Consistent with previous studies^20^, primary MM exhibited significantly fewer clonal mutations but substantially greater heterogeneity, purity, ploidy, and aneuploidy than CM (**Fig. 1b** and **Supplementary Table 4**), and these differences were preserved across anatomic sites (**Extended Data Fig. 1e**). Whereas *BRAF* mutations predominated the CM primary tumors, *SF3B1* was the most frequently altered gene in primary MM (**Fig. 1c**, **Extended Data Fig. 2a** and **Supplementary Table 4**), as previously reported^21^. Mutational signature analysis further distinguished MM from CM. The UV-associated mutational signature, single-base substitution (SBS) 7, predominated in CM, whereas an age-associated signature, SBS1, was the dominant mutational process in MM (**Fig. 1d** and **Supplementary Table 4**). Copy number alteration analysis showed chromosomal arm-level losses were more frequent in MM than in CM (**Fig. 1e**). All chromosomal arms that reached significance showed more frequent loss in MM, with the largest differences on 16p (52% versus 10%), 3q (48% versus 10%), 6q (83% versus 48%), 16q (65% versus 38%) and 8p (64% versus 33%). In contrast, no chromosome arms differed significantly in gain frequency (**Extended Data Fig. 2b**), and the genome-wide frequency profiles indicate that the divergence between the two primary tumor types is driven by loss rather than gain (**Extended Data Fig. 2c** and **Supplementary Table 4**).

Transcriptomic profiling further revealed that primary MM tumors were enriched for proliferation (Mitotic Spindle)-, dedifferentiation (e.g., Mesenchymal like, Wouters. MES-like)-, and epithelial–mesenchymal transition (EMT)-associated programs, whereas primary CM retained stronger melanocytic lineage signatures (e.g., Wouters. Melanocytic) (**Fig. 1f** and **Supplementary Table 5**). These transcriptional differences were further supported by Orion imaging, which demonstrated a significantly higher proportion of proliferating SOX10⁺KI67⁺ tumor cells in primary MM than in primary CM (**Fig. 1g, h**). Taken together, these findings demonstrate that the intrinsic biological divergence between MM and CM is already established at the primary tumor stage, before metastatic progression or therapeutic intervention.

### MM has distinct genetic subtypes

Though genomic drivers of MM have been previously evaluated^20^, genomic classification of MM, i.e. distinct subgroups of MM with mutually exclusive genomic drivers, has not been accomplished to date. Because primary and metastatic tumors showed no significant difference in genomic features **(Extended Data Fig. 2d)**, all samples were combined to increase the number of tumors available for subtype identification. To identify genetic subtypes, we analyzed MM targeted sequencing data from DFCI (n=86)^22^ **(Supplementary Table 2)** and validated our findings in a non-overlapping cohort from the AACR Project GENIE (n=253)^40^ (**Fig. 2a, Extended Data Fig. 3a, and Supplementary Table 3**). These cohorts exhibited similar stage and anatomic site distributions (**Fig. 2b, Extended Data Fig. 3b,c**). For genomic classification, we focused on 23 previously reported MM genetic alterations (15 oncogenes and 8 tumor suppressors)^20^. We examined genes altered in more than 10% of patients in both cohorts and identified only one gene not included in our list, CDKN2B, which is co-located with CDKN2A (**Extended Data Fig. 3d**), and thus subsequently utilized the 23-gene panel for subtype analyses.

**Fig. 2:**
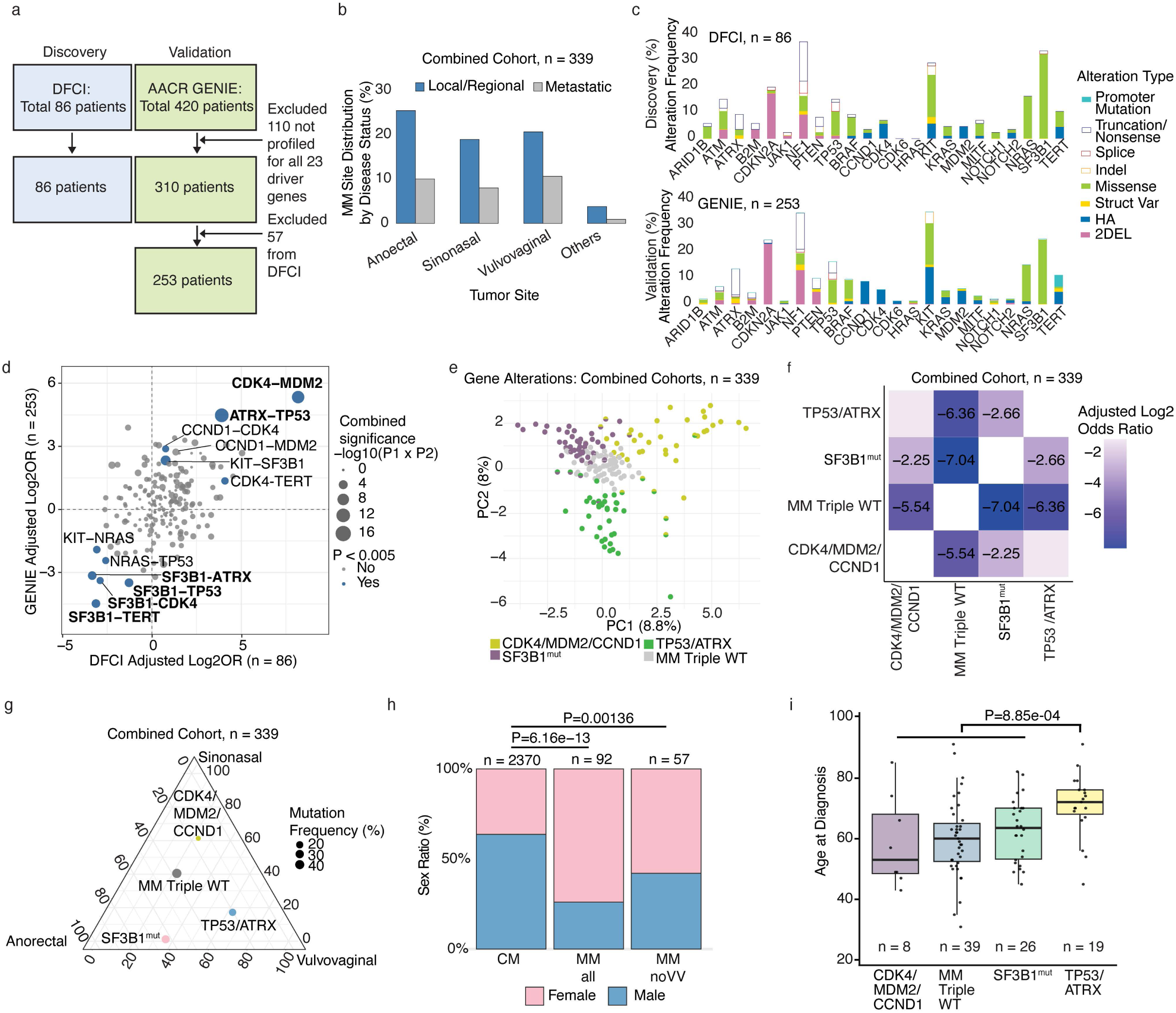
Distinct genetic subtypes of mucosal melanoma. **a**, Targeted exome patient data schematic. **b**, Site distribution and disease status for combined cohort. **c**, Gene alteration frequencies in DFCI (top) and GENIE (bottom) cohorts. HA = high amplification, Indel = insertions and deletions, Struct Var = structural variations, 2DEL = two copy deletion. **d**, Gene-pair association scatter plot comparing DFCI and GENIE cohorts; each point shows log2(OR) in DFCI versus GENIE, with significant pairs highlighted in blue. **e**, Principal component analysis of gene alteration data in the combined cohort. Each point represents one patient colored by key alterations. **f**, Heatmap of mutual exclusivity analysis among MM genetic subtypes (Fisher’s exact test). The heatmap is colored as per adjusted Log2(OR) and corresponding numerical values are depicted in each block. **g**, Ternary plot of site-specific MM genetic subtype distribution across primary tumor sites in the combined cohort. The size of dots represents the number of patients. **h**, Sex distribution in CM (AACR GENIE cohort) and MM (AACR GENIE cohort + DFCI data), including analysis after exclusion of vulvovaginal MM cases (Fisher’s exact test). **i**, Age distribution across MM subtypes. Groups were compared using a two-sided Fisher’s exact test.

Alteration types and frequencies of the 23 genes in both cohorts were remarkably consistent (**Fig. 2c, Extended Data Fig. 3e,f**). NF1 (loss/mutation), KIT (mutation/copy number gain), SF3B1 (point mutation), and CDKN2A (deletion) were the most frequent alterations. SF3B1 had the most point mutations, and 98.9% of the SF3B1 mutations were found at residue 625 (90/91), despite the rarity of point mutations in MM (**Fig. 2c**), suggesting either a distinct mutational mechanism or strong selective pressure for these mutations.

To evaluate the global genetic architecture of MM, we examined co-occurrence and mutual exclusivity of driver alterations in the 23-gene panel in both cohorts (**Fig. 2d, Extended Data Fig. 3g,h,** and **Extended Data Table 1**). We found that amplifications in CDK4 and MDM2, adjacent genes on chromosome 12, were strongly co-occurring in both cohorts (DFCI: OR = 289.8, Fisher’s exact *P* = 7.06 × 10^−6^; GENIE: OR = 40.5, Fisher’s exact *P* = 4.4 × 10^−8^). We also found that loss/mutation of TP53 and ATRX were strongly co-occurring in both cohorts (DFCI: OR = 14.9, Fisher’s exact *P* = 1.0 × 10^−3^; GENIE: OR = 22.5, Fisher’s exact *P* = 9.34 × 10^−14^), an association seen in other cancers^43–45^. Further, in both cohorts we found that SF3B1 mutations were mutually exclusive with CDK4 amplifications (DFCI: OR = 0.13, Fisher’s exact *P* = 9.0 × 10^−2^; GENIE: OR = 0.09, Fisher’s exact *P* = 2.4 × 10^−2^) and TP53 loss/mutation (DFCI: OR = 0.4, Fisher’s exact *P* = 3.2 × 10^−1^; GENIE: OR = 0.09, Fisher’s exact *P* = 5.5 × 10^−5^), suggesting the presence of distinct genetic subgroups. To test whether mutation associations were confounded by differences in tumor mutational burden (TMB), we estimated empirical p-values for observed gene co-occurrence and mutual exclusivity compared to a TMB-controlled null distribution in the larger GENIE cohort^40^ (**Extended Data Fig. 3i and Supplementary Table 6**). This analysis (limited methodologically to point mutations, hence excluding CDK4/MDM2 amplifications) confirmed significant co-occurrence between TP53 and ATRX mutations (p < 0.001), mutual exclusivity between SF3B1 and TP53 (p < 0.001), and verified a significant association between SF3B1 and KIT mutations (p < 0.001).

To map these gene-gene associations to patients, we performed a principal component analysis on the 23-gene alterations across the combined cohort. We observed spatially distinct groups (on the PC1/PC2 axes) defined by SF3B1^R625^ alterations (26.8%), TP53/ATRX alterations (20.3%), and CDK4/MDM2 amplifications (*n* = 8%) (**Fig. 2e**). We further observed that CCND1 amplification co-occurred with both CDK4 (GENIE: OR = 7.3, Fisher’s exact *P* = 3.7 × 10^−3^) and MDM2 (GENIE: OR = 6.6, Fisher’s exact *P* = 5.2 × 10^−3^) in the larger GENIE cohort and spatially co-occurred with the CDK4/MDM2 group in the PCA visualization. As CCND1 and CDK4 both represent amplification of cell cycle genes^46^, we included CCND1 amplification with CDK4/MDM2 amplification to define a CDK4/MDM2/CCND1 amplified subtype (13.5%) characterized by dysregulated cell-cycle progression. Patients were then assigned to subtypes based on the presence of at least one defining alteration within a subtype.

We designated the remaining MM tumors not assigned to any subtype as MM Triple Wild Type (WT). We found that the MM Triple WT group was enriched for NRAS, TERT, and BRAF alterations (**Extended Data Fig. 3j**). Notably, these alterations are more common in CM^25^, suggesting that this group of MMs may be more CM-like. Finally, we confirmed that these genetic subgroups were mutually exclusive in both the DFCI and GENIE cohorts (**Extended Data Fig. 3k–l;** combined cohort shown in **Fig. 2f**). Taken tother, these analyses defined four distinct genetic subtypes of MM: SF3B1-mutant; TP53/ATRX-mutant; CDK4/MDM2/CCND1 amplified; and MM Triple WT.

### Genomic and clinical correlates of genomic subtypes

We next analyzed the anatomic site distribution and the total cumulative alteration frequency for each subtype from the combined cohort. As previously reported, SF3B1^R625^ occurred almost exclusively in anorectal and vulvovaginal MM (OR = 56.4, Fisher’s exact *P* = 1.3 × 10^−14^, 97%)^20,47,48^. We found that the CDK4/MDM2/CCND1 subtype occurred more commonly in sinonasal MM (OR = 5.15, Fisher’s exact *P* = 1.1 × 10^−6^, 59%), while TP53/ATRX occurred more commonly in vulvovaginal MM (OR = 2.21, Fisher’s exact *P* = 0.03) (**Fig. 2g**). MM Triple WT tumors were not associated with any primary site. MM from vulvovaginal sites (Kolmogorov-Smirnov, *P* = 0.028, **Extended Figure Fig. 4a**) and the CDK4/MDM2/CCND1 subtype (Kolmogorov-Smirnov, *P* = 0.001, **Extended Data Fig. 4b**) had the most genetic alterations.

To identify the clinical characteristics of the MM subtypes, we analyzed data for 97 DFCI patients (86 patients and 11 newly available patients). There was no significant association between the MM subtypes and cancer stage at diagnosis (**Extended Data Fig. 4c**), or with sex. Notably, all MMs showed a higher proportion of female patients than CM, even after exclusion of vulvovaginal MMs (Fisher’s exact *P* = 0.001, **Fig. 2h**), validating a prior report^49^. Patients with the TP53/ATRX subtype were significantly older (median age TP53/ATRX 72, all other groups 61, Wilcoxon rank-sum *P* = 8.85 × 10^−4^, **Fig. 2i**), and fewer received adjuvant therapy after surgery (*P* = 0.047, **Extended Fig. 4d**). Interestingly, the small subset of patients with complex subtypes (5.15% of MM patients in our cohort had alterations in genes from two or more subtypes) had a significantly higher frequency of prior malignancies before melanoma diagnosis (*P* = 0.01, **Extended Data Fig. 4e**).

We then evaluated treatment patterns and clinical outcomes across the MM subtypes. Across subtypes, we found that adjuvant medical therapy was not associated with improved recurrence-free survival compared to those who did not receive adj therapy (Log-rank *P* = 0.93, **Extended Fig. 4f**). Recurrence locations (local, nodal, or distance) were similar across subtypes (**Extended Data Fig. 4g**). Unsurprisingly, patients with metastatic disease had higher rates of depression and anxiety than those without metastatic disease (Fisher’s exact *P* = 0.035, **Extended Fig. 4h**). Overall survival, chance of developing brain metastases, first-line systemic therapy at metastasis, and first-line therapy outcomes were not statistically significantly different across subtypes (**Extended Data Fig. 4i-l**). Overall, we find that under current treatment regimens, treatment-related clinical outcomes are similar across MM genetic subtypes.

### Enrichment of CD163^+^ TAMs distinguishes the immune microenvironment of MM

To characterize the immune microenvironment of MM, we performed multiplex Orion imaging using markers for tumor cells (SOX10 and MART1), stromal structures (CD31 and αSMA), and major immune populations, including T cells (CD3E, CD4, CD8A and FOXP3), B cells (CD20), myeloid cells (CD11c), and CD163⁺ TAMs. Primary MM showed lower overall CD8^+^ T-cell infiltration than primary CM, with marked heterogeneity among primary MM tumors (**Fig. 3a**). We noticed that some MM primaries were immune-cold, whereas others were enriched with CD8^+^ T cells and CD163^+^ TAMs. We quantified immune infiltration within tumor compartments using SOX10⁺ tumor boundaries validated by matched H&E on the same slide (**Fig. 3b** and **Extended Data Fig. 5a, b**). Compared with primary CM, primary MM contained significantly fewer tumor-infiltrating T cells—including CD8⁺ T cells, CD4⁺ T cells, Tregs, and CD11c⁺ myeloid cells— but significantly greater infiltration of CD163⁺ TAMs (**Fig. 3c**, **Extended Data Fig. 5c** and **Supplementary Table 7**). These findings identify decreased T cell and increased CD163⁺ TAM enrichment as distinct features of the MM immune microenvironment.

**Fig. 3:**
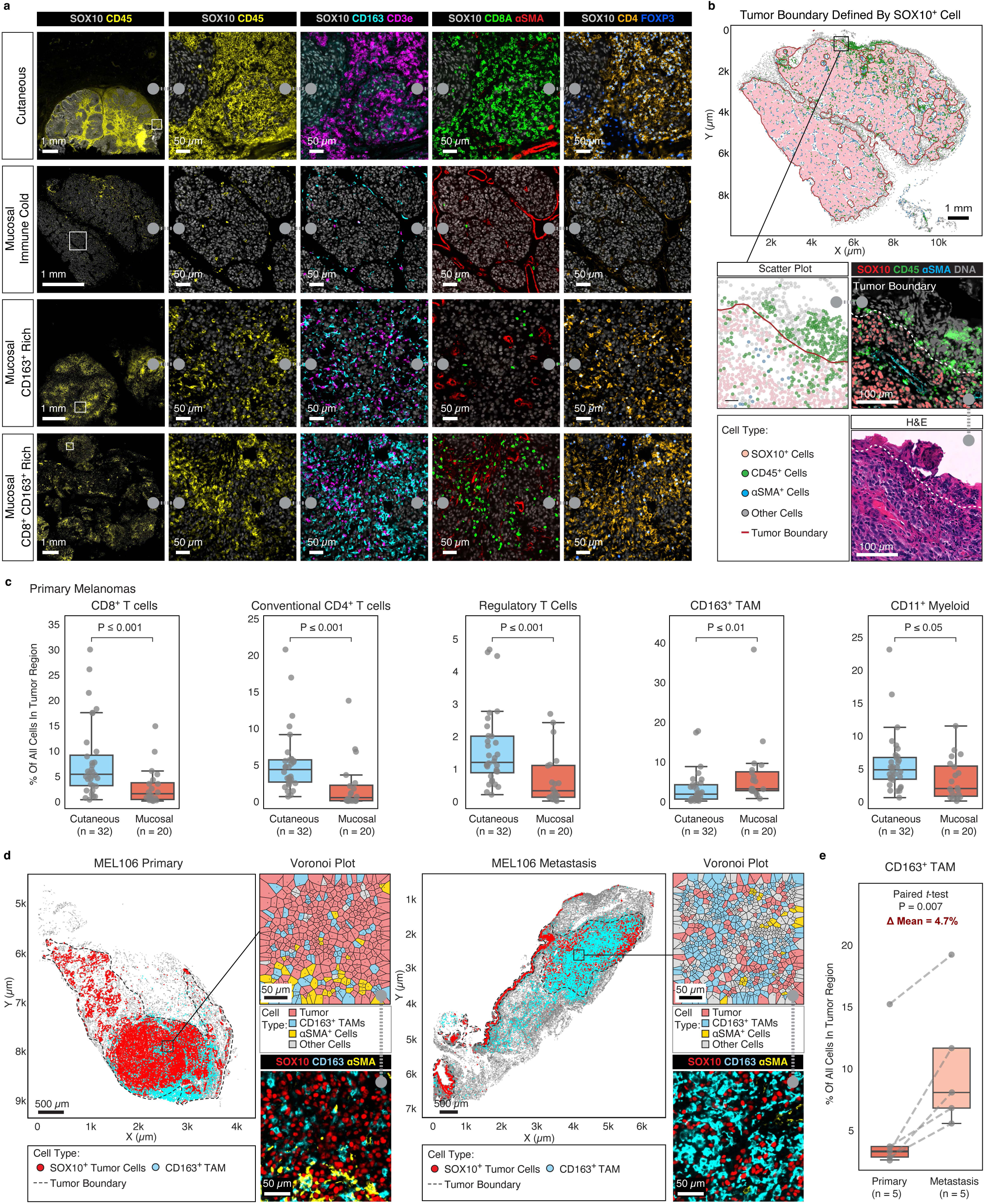
CD163⁺ TAMs define the immune microenvironment of MM. **a**, Representative Orion multiplexed immunofluorescence images illustrating the spectrum of immune infiltration in primary CM and primary MM, including immune-cold, CD163⁺-rich, and CD8⁺/CD163⁺-rich tumors. Tumor cells were identified by SOX10, with immune cell populations visualized using CD45, CD163, CD11c, CD3E, CD8A, CD4, and FOXP3. Higher-magnification views of the boxed regions are shown. Scale bars, 1 mm and 50 μm. **b**, Schematic illustrating the definition of tumor compartment and quantification of tumor-infiltrating immune cells. Tumor boundaries were defined using SOX10 expression and validated by matched H&E staining on the same tissue section. Tumor, tumor border, and stromal compartments were subsequently annotated for downstream spatial analyses. Scale bars, 1 mm and 100 μm. **c**, Quantification of tumor-infiltrating immune cell populations within the tumor compartment in primary CM (*n* = 32) and primary MM (*n* = 20). Shown are the proportions of CD8⁺ T cells, conventional CD4⁺ T cells, regulatory T cells, CD163⁺ TAMs, and CD11c⁺ myeloid cells within the tumor compartment. Box plots show the median and interquartile range (IQR), with whiskers extending to 1.5 ’IQR and individual tumors shown as points. Groups were compared using two-sided Mann-Whitney *U* test. Each point represents one tumor. **d**, Representative spatial maps and corresponding Voronoi tessellations illustrating the distribution of SOX10⁺ tumor cells and CD163⁺ TAMs in matched primary and metastatic MM tumors from the same patient (MEL106). Insets show higher-magnification regions with corresponding Orion multiplex immunofluorescence images stained for SOX10 (red), CD163 (cyan), and aSMA (yellow). Tumor boundaries are indicated by dashed lines. Scale bars, 500 μm and 50 μm. **e**, Paired comparison of tumor-infiltrating CD163⁺ TAM abundance between matched primary and metastatic MM samples. Dashed lines connect paired samples. Statistical significance was determined using the paired student *t*-test.

To determine whether CD163⁺ TAM accumulation accompanies disease progression, we compared treatment-naïve primary and metastatic MM tumors. Although CD163⁺ TAM abundance did not differ significantly between unmatched primary and metastatic tumors (**Extended Data Fig. 5d** and **Supplementary Table 8**), a paired analyses of treatment-naïve primary and metastatic tumors from the same patients (n = 5 patients) demonstrated a significantly increased CD163⁺ TAM infiltration in metastatic lesions (paired T-test *P* = 0.007, **Fig. 3d,e**, **Extended Data Fig. 5e**, **Extended Data Fig. 6a,b** and **Supplementary Table 8**), suggesting that this macrophage subset may increase during disease progression.

### Genomic instability and TAM-associated immunosuppressive niche characterize immunotherapy resistance in MM

To investigate the intrinsic and extrinsic determinants of immunotherapy response in MM, we stratified patients by clinical outcome following first-line ICB (**Fig. 4a**). Median progression-free survival (PFS) to ICB in MM patients is < 6 months^11^. However, a small subset of MM patients experiences better outcomes. We defined durable responders (DR) as patients with PFS ≥ 12 months and rapid progressors (RP) as patients with PFS ≤ 6 months. Patients with PFS between 6 and 12 months were classified as an intermediate (INT) group (**Fig. 4a** and **Supplementary Table 9**).

**Fig. 4:**
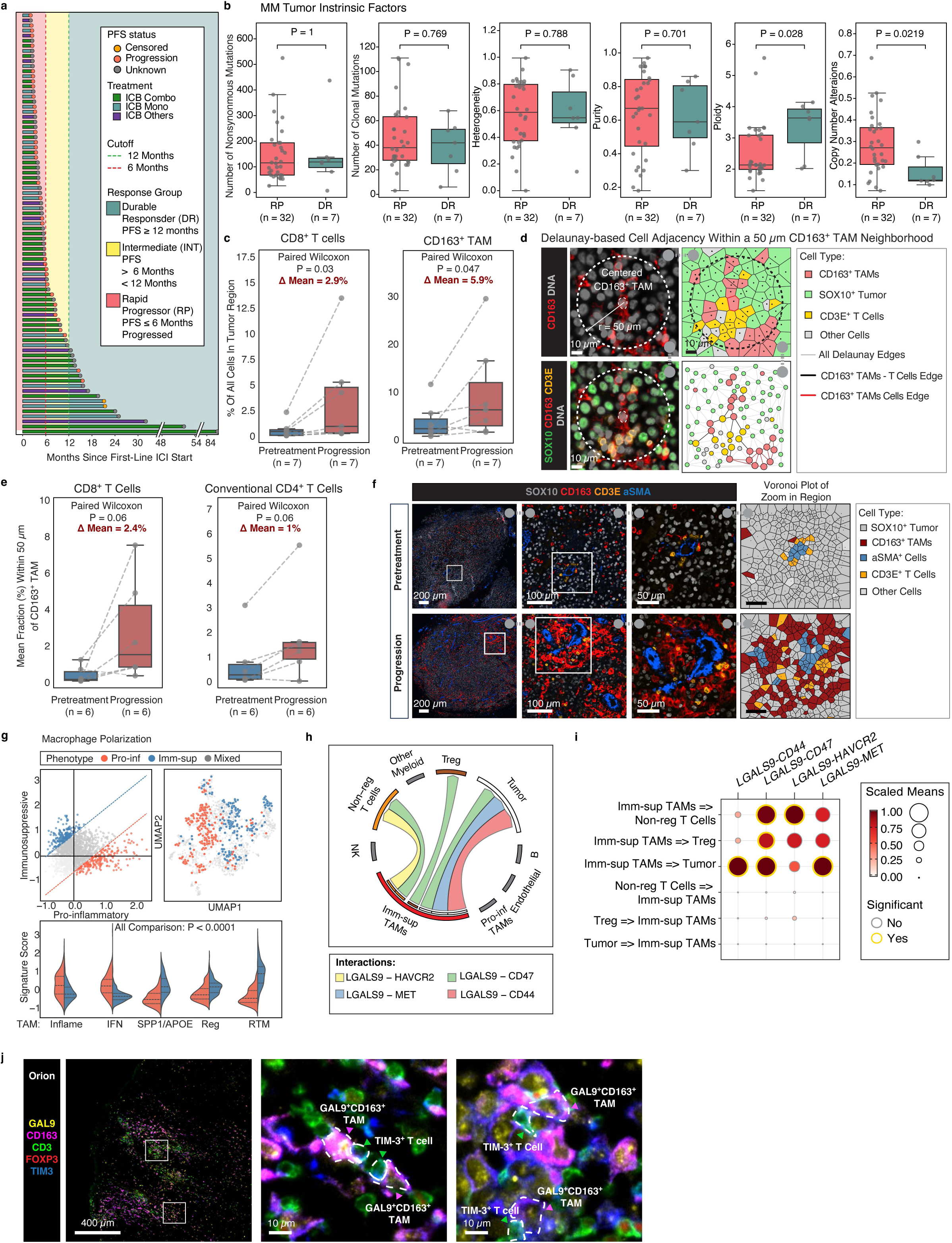
Genomic instability and a CD163⁺ TAM-mediated immunosuppressive niche are associated with immunotherapy resistance in mucosal melanoma. **a,** Patient stratification according to PFS following first-line ICB. Each bar represents one patient, with bar length indicating PFS. Bar color denotes the first-line regimen: ICB combo (green), ICB mono (light blue), and other ICB-containing regimen (purple); the terminal point denotes PFS status: progression (red), censored (orange), and unknown (grey). Background shading indicates defined response groups: DR (teal), INT (yellow), and RP (pink). Dashed lines mark the 6-month (red) and 12-month (green) cutoffs. Axis breaks are shown beyond 48 months. **b,** Comparison of genomic features between DR (*n* = 7) and RP (*n* = 32) pretreatment tumors. Shown are nonsynonymous mutation burden, clonal mutation burden, intratumoral heterogeneity, tumor purity, ploidy, and the proportion of copy-number alterations derived from WES. Box plots show the median and interquartile range (IQR), with whiskers extending to 1.5 × IQR and individual tumors shown as points. Groups were compared using two-sided Mann-Whitney *U* tests; significant *P* values (*P* < 0.05) are shown. **c,** Paired comparison of tumor-infiltrating CD8⁺ T cells and CD163⁺ TAMs between pretreatment and matched post-progression MM tumors (*n* = 7 pairs). Dashed lines connect paired samples. Groups were compared using two-sided Wilcoxon signed-rank tests. **d,** Schematic of the neighborhood analysis workflow. CD163⁺ TAMs were defined as the central cell population, and neighboring cells within a 50-μm radius were identified using Delaunay-based cell adjacency to quantify local cellular composition. Scale bars, 10 μm. **e,** Neighborhood analysis comparing the mean fraction of neighboring CD8⁺ and conventional CD4⁺ T cells surrounding CD163⁺ TAMs in paired pretreatment and progression tumors (*n* = 6 pairs). One SOX10-negative pair was excluded; tumor regions were annotated by H&E, but individual tumor cells could not be identified for neighborhood analysis. Dashed lines connect paired samples. Groups were compared using two-sided Wilcoxon signed-rank tests. **f,** Representative Orion multiplexed immunofluorescence images of matched pretreatment and progression tumors showing enrichment of CD163⁺ TAMs in regions containing T cells. Corresponding spatial maps illustrate the distribution of CD163⁺ TAMs (red) and neighboring CD3E⁺ T cells (orange) identified within the CD163⁺ TAM neighborhood. Scale bars, 200 μm, 100 μm, and 50 μm. **g**, Macrophage polarization analysis of scRNA-seq data. Left, macrophage polarization based on pro-inflammatory and immunosuppressive gene signatures. Top right, UMAP visualization of polarized macrophage subsets. Bottom, comparison of published macrophage gene signatures across polarized macrophage populations. **h**, Circos plot summarizing inferred ligand–receptor interactions between immunosuppressive CD163⁺ TAMs and neighboring cell populations. **i**, Dot plot showing *LGALS9*-mediated ligand–receptor interactions between immunosuppressive CD163⁺ TAMs and other cell populations. Dot size represents the scaled mean interaction strength, and color indicates relative ligand expression. Red outlines denote statistically significant interactions (permutation *P* < 0.05). **j**, Representative Orion multiplexed immunofluorescence images from progression tumors showing GAL9 (yellow), CD163 (magenta), CD3 (green), FOXP3 (red), and TIM-3 (cyan). Insets show higher-magnification views illustrating GAL9⁺CD163⁺ TAMs in close proximity to TIM-3⁺ T cells. Scale bars, 400 μm and 10 μm.

We analyzed pre-treatment tumors with WES and bulk RNA sequencing to identify tumor-intrinsic features associated with response. There were no significant differences in mutation burden or intratumoral genomic heterogeneity between DR and RP groups. However, RP tumors exhibited significantly lower ploidy but a higher proportion of copy-number alterations than DR tumors (*P* = 0.028 and *P* = 0.022, respectively **Fig. 4b, Extended Data Fig. 7a** and **Supplementary Table 10**). Transcriptomic analysis further showed a trend toward higher dedifferentiation- and EMT-associated signature scores in RP tumors compared with DR tumors (**Extended Data Fig. 7b**). Baseline immune composition was then evaluated using Orion imaging of pretreatment tumor samples. No significant differences in the abundance of major immune cell populations were observed between RP and DR tumors (**Extended Data Fig. 7c** and **Supplementary Table 11**).

To identify features associated with resistance to ICB, we compared pretreatment and post-progression MM tumors using both unpaired and paired same-patient analyses. Both comparisons showed a consistent increase in clonal mutation burden at progression (**Extended Data Fig. 7d**, **Extended Data Fig. 8a** and **Supplementary Table 12**), suggesting clonal selection pressures. In the TME, we found that CD163^+^ TAMs and CD8^+^ T cells were significantly enriched in progression tumors in paired samples (n = 7 patients; **Fig. 4c** and **Extended Data Fig. 8b, c** and **Supplementary Table 13**). To determine the spatial association of these populations, we performed neighborhood analysis using CD163⁺ TAMs as the central cell population and quantified neighboring cell types within a 50-μm radius (**Fig. 4d**). Compared with pretreatment tumors, progressing tumors exhibited a trend toward increased enrichment of neighboring non-regulatory T cells (CD8⁺ and CD4⁺) surrounding CD163⁺ TAMs (*P* = 0.06, **Fig. 4e,f** and **Extended Data Fig. 8d**).

To further characterize the interactions between CD163⁺ TAMs and neighboring T cells, we analyzed scRNA-seq data from eleven MM tumors collected from patients who progressed following immunotherapy^22^. A total of 1720 macrophages were identified. Polarization analysis classified these populations into pro-inflammatory and immunosuppressive subsets (see Methods), which exhibited distinct transcriptional profiles (**Fig. 4g, top**). Confirming this characterization, we found that immunosuppressive TAMs were significantly enriched for SPP1/APOE, regulatory macrophage (Reg), and resident tissue macrophage (RTM) programs relative to pro-inflammatory TAMs (**Fig. 4g, bottom, Extended Data Fig. 8e** and **Supplementary Table 14**).

To evaluate the function of the immunosuppressive macrophage population, we inferred their ligand-receptor interactions using single cell RNAseq. Among the top statistically significant predicted interactions, we found *LGALS9*-mediated interactions between immunosuppressive TAMs and tumor cells (*MET*, *CD47*, and *CD44*), as well as non-regulatory T cells (*HAVCR2*, also known as TIM3) (**Fig. 4h,I** and **Supplementary Table 15**). To validate the corresponding Galectin-9(GAL9)-TIM3 protein interaction with an orthogonal assay, we performed Orion imaging on matched progressing tumors from the scRNA-seq cohort. Consistent with the ligand–receptor inference, GAL9⁺CD163⁺ TAMs were spatially localized in close proximity to TIM-3⁺ T cells within the TME (**Fig. 4j**). Together, these complementary transcriptomic and spatial analyses identified the GAL9–TIM3 axis as a prominent feature of the immunosuppressive macrophage niche in MM following immunotherapy progression.

## DISCUSSION

In this study, we integrate clinical, genomic, transcriptional, and spatial profiling across a multi-institutional cohort to provide a comprehensive molecular framework for MM. We demonstrate that at the primary stage before metastasis or treatment exposure, MM already has distinct biology from CM, independent of the site of origin. With analysis of the largest cohorts of genomically characterized MM to date, we demonstrate the first genomic classification of MM at scale. Finally, we demonstrate that, distinct from CM, MM exhibits a macrophage-enriched immunosuppressive microenvironment whose interactions may contribute to MM’s poor response to ICB. Together, these findings provide insight into the biology underpinning MM’s aggressive clinical phenotypes, therapy resistance, and provide a framework to advance therapeutic approaches based on distinct genomic subtypes and specific immunosuppressive interactions in the microenvironment.

We observe biological differences between MM and CM established early during tumorigenesis, before metastatic progression or treatment exposure. Compared with primary CM, primary MM accumulated fewer clonal mutations but exhibited substantially greater genomic instability, characterized by increased intratumoral heterogeneity, ploidy, and aneuploidy, which has been described previously^20,50^. Transcriptionally, primary MM was enriched for proliferative, dedifferentiated, and EMT programs, whereas CM retained stronger melanocytic lineage programs, consistent with MM’s aggressive clinical behavior. Together, these findings indicate that MM has a different cell intrinsic transcriptional state and follows a fundamentally different evolutionary trajectory from CM.

We identify four mutually exclusive genetic subtypes of MM: *CDK4/MDM2/CCND1*-amplified, *TP53/ATRX*-altered, *SF3B1*^R625^–mutant, and MM Triple WT. Some of these co-occurrences and clinical correlations have been previously reported, including CDK4/MDM2 amplifications^20,51^, TP53 loss/mutation with ATRX alteration in MM^20^ and other cancers^43–45^. Utilizing the largest genomically characterized collection of MM to date, we were able to comprehensively analyze co-occurring and mutually exclusive genomic alterations to develop the first genomic classification of MM.

The molecular framework established in this study provides opportunities for precision approaches for MM. Distinct driver alterations in each genomic subtype implicate different biological pathways and are associated with some distinct clinical features and presentations. For example, SF3B1-mutated primaries are concentrated in anorectal and vulvovaginal locations^20,47,48^ and TP53/ATRX-altered MMs are enriched in older patients, which is consistent with reports that TP53 mutations are also linked to older age in CM^52^ and in advanced solid tumors more broadly^53^. Despite this, we did not observe significant differences in PFS, overall survival, recurrence-free survival after immune therapy, or response to first-line immunotherapy across genomic subtypes. Rather than suggesting biological equivalence, this likely reflects that patients currently receive subtype-agnostic treatments that do not exploit underlying molecular differences. For future directions, our findings suggest that targeted approaches for MM genetic subtypes may be a fruitful direction. For example, CDK4/MDM2/CCND1-amplified tumors may represent candidates for CDK4/6-directed therapies, whereas the distinct biology of SF3B1^R625^– mutant tumors will require different targeted therapies. Our findings further provide the rationale for and guide the development of subtype-specific experimental MM models^22^ that can be used to dissect mechanisms and develop therapeutics. Broadly, this framework may provide a foundation for biologically informed therapeutic strategies in this challenging disease.

Beyond tumor-intrinsic features, our integrative spatial and single-cell analyses provide new insights into the immune landscape of MM. Previous studies have emphasized the relatively low abundance of TILs in MM compared with CM, leading to the concept of MM as an immunologically "cold" tumor^54^. Indeed, we confirm lower TILs in MM (even at primary stages), but we identify significant heterogeneity in immune infiltration. Our findings refine this model. We observe consistent enrichment of CD163^+^ TAMs relative to CM, even at the primary stage of disease. CD163 is widely used as a marker of TAMs and has been associated with immunosuppressive TME across multiple cancer types^55–58^. Consistent with this concept, our longitudinal analyses suggested that CD163^+^ TAMs become increasingly prominent during acquired resistance to ICB in a subset of patients. The integration of spatial neighborhood analysis, macrophage polarization, ligand–receptor inference, and multiplex imaging collectively implicates interactions between immunosuppressive macrophages and T cells through the GAL9–TIM3 axis. These observations support a model in which macrophage-mediated immune suppression contributes to immune escape in MM, rather than immune evasion arising from reduced T-cell infiltration. GAL9 is a secreted immunoregulatory lectin released by macrophages into the TME, where it binds the inhibitory receptor TIM-3 expressed on activated and exhausted T cells. Engagement of the GAL9–TIM3 axis has been shown to suppress T-cell effector function, promote T-cell exhaustion and, in some contexts, induce T cell apoptosis, thereby dampening antitumor immunity^59–61^. Notably, this pathway has also been implicated in resistance to ICB in CM and other solid tumors^61,62^, and a first-in-class anti-Gal9 monoclonal antibody (LYT-200) is currently undergoing phase 1/2 clinical trial as monotherapy and in combination with PD-1 blockade in patients with advanced solid tumors (NCT04666688). Thus, GAL9 blockade may represent a promising new therapeutic strategy for MM.

Our analysis of the rare MM patients with DR to ICB suggests that these tumors have distinct underlying tumor-intrinsic biology. Genomically, they had higher ploidy but a lower proportion of copy-number alterations. Transcriptionally, DR exhibited a more differentiated tumor program, consistent with previous reports associating dedifferentiation and neural-crest-like states with immunotherapy resistance in CM^63–65^. Interestingly, we did not observe significant differences in the immune microenvironment between pre-treatment and post-progression MM tumors following ICB. However, compared with matched pre-treatment samples, post-ICB resistant samples showed an increased number of clonal mutations, suggesting clonal selection and outgrowth during immunotherapy, as well as TAM enrichment as previously discussed., Overall, our data suggests that both tumor-intrinsic and immune programs are associated with ICB response and resistance.

### Limitations

Although this represents one of the largest multi-omic studies of MM to date, additional focused questions would be better answered in future studies. We observe association of anatomical site with some genomic features and subtype, but how biology may differ both in tumorigenesis and subsequent tumor evolution across sites is unclear and we are underpowered to further probe more subtle anatomic specific differences. Our data demonstrates the enrichment of immunosuppressive TAMs in MM and nominates GAL9-TIM-3 interactions in tumor aggressiveness and treatment resistance. This interaction should be functionally validated and should be confirmed in an independent cohort.

In summary, our study demonstrates that MM is biologically distinct from CM, with unique tumor programs and immune interactions evident from the earliest stages that are associated with its aggressive clinical phenotype and resistance to standard CM therapies. We define a molecular classification of MM to enable precision-based therapeutic approaches and identify macrophage-mediated immunosuppression as a candidate mechanism of ICB resistance. Together, these findings provide a biological framework for future site-informed diagnosis, mechanistic investigation, and precision therapeutic development in this rare and aggressive melanoma subtype.

## Supporting information

Extended_Data_Table_1 Cooccurence and mutual exclusivity in DFCI and GENIE cohort

## ACKNOWLEDGEMENT

The authors would like to acknowledge the DFCI Oncology Data Retrieval System (OncDRS) for the aggregation, management, and delivery of genomic research data used in this project.

## AUTHOR CONTRIBUTION STATEMENTS

Y.S., P.S., M.L.I., D.L. and G.M.B. conceived and designed the study. D.L., G.M.B., M.L.I. and P.K.S. supervised the work. Y.S. and P.S. performed the primary data analysis. Y.S. performed the whole-exome, copy-number, mutational signature, transcriptomic and Orion spatial analyses. P.S. performed the targeted sequencing analysis. J.-R.L. developed and performed the multiplexed Orion imaging, image processing and preliminary image analysis, with input from T.V., R.P., M.L.L., Y.D.L. and S.M.P., D.A., M.P.M. and L.R.A. provided clinical samples and clinical annotation. G.T. contributed computational analysis of whole-exome sequencing and transcriptomic data. N.A. and R.B. contributed to the permutation analysis. J.C. contributed to the scRNA-seq analysis. M.R. contributed to figure preparation. C.G.L. performed histopathological review. D.E.F., R.B. and E.I.B. provided scientific input and interpretation. Y.S., P.S., M.L.I. and D.L. wrote the manuscript with input from G.M.B. All authors reviewed and approved the final manuscript.

## DATA AND SOFTWARE AVAILABILITY

Software environments and code to reproduce all analyses and generate figures is available on GitHub (https://github.com/davidliu-lab/mucosal_melanoma_atlas_manuscript_code_2026). Additional reasonable requests for code will be promptly reviewed by the senior authors to verify whether the request is subject to any intellectual property or confidentiality obligations and will be shared to the extent permissible by these obligations. All image data supporting this study are available via amazon S3. Instructions for accessing imaging data and a data index page can be found on GitHub (https://github.com/labsyspharm/2026-Shi-Genomic-classification-and-immune-profiling-define-mucosal-melanoma), which has been archived on Zenodo (https://doi.org/10.5281/zenodo.23423190). The WES and bulkRNA will be released on dbGap.

## METHODS

### Patient cohort and specimen collection

Melanoma specimens were obtained from patients who provided written informed consent under Institutional Review Board (IRB)-approved biospecimen collection protocols at the Dana-Farber/Harvard Cancer Center (DF/HCC; protocol 05-042 for DFCI patients and 11-181 for MGH patients). Analyses of residual clinical specimens were performed under DF/HCC secondary-use protocols 20-171 and 23-563 with an IRB-approved waivers of documentation of informed consent under 45 CFR 46.117(c). All studies were conducted in accordance with the Declaration of Helsinki and approved by the DF/HCC IRB.

### Clinical outcome classification

DR and RP were defined by PFS from initiation of first-line ICB (see Results), applied identically across the DFCI and MGH cohorts. Because this definition is based on time to progression rather than radiographic BOR category, sustained stable disease is captured within the DR group when the corresponding PFS criterion is met. Outcome groups were derived by identical criteria in both cohorts. PFS was defined as the interval from the first dose of first-line ICB to radiographic or clinical progression, or to death from any cause; patients without a progression event were censored at the date of last imaging assessment. For the analysis of features associated with immunotherapy response, we excluded patients with PFS < 2 months as patients with immediate clinical progression on treatment where no treatment was likely to alter the course of disease.

### Whole exome sequencing

WES for samples from MGH and DFCI cohorts was performed at the Broad Institute Genomic Platform on 133 tumor-matched normal blood sample pairs from 84 patients, using a previously described protocol^66^. Briefly, 150–500 ng of genomic DNA (gDNA) was extracted using QIAGEN AIIPrep DNA/RNA Mini Kit (cat# 80204). DNA was fragmented by acoustic shearing, followed by size selection to achieve a library insert size distribution of 300– 650 ng. Libraries were prepared using the KAPA HyperPrep Kit according to the manufacturer’s instructions and subsequently quantified and normalized using PicoGreen to ensure equal concentrations. Adapter ligation was performed using the Illumina TruSeq DNA Exome Kit according to the manufacturer’s instructions. Libraries were sequenced using an HiSeq2500 with paired end 76bp reads, followed by analysis with RTA v.1.12.4.2.

### WES quality control and variant calling

Initial WES data processing and analysis were performed using a customized version of the Getz Labs WES analysis pipeline at the Broad Institute. After alignment from the Broad Picard Pipeline, BAM files were uploaded into the Terra infrastructure (https://app.terra.bio), which managed intermediate analysis files executed by analysis pipelines. Samples were required to meet the following thresholds: mean target coverage > 50× for tumor samples and >30× for matched normal samples, cross-sample contamination (ContEst) <5%, tumor purity ≥15%, and DeTiN-estimated tumor-in-normal (TiN) ≤ 20%. A power-based filter integrating sequencing coverage and tumor purity was applied as previously described to ensure ≥80% power for detecting clonal mutations^50,67^. samples were excluded because of low tumor purity.

MuTect2 (RRID:SCR_026692)^68^ was used to identify somatic single nucleotide variants (SNVs) in targeted exons, with computational filtering to remove artifacts arising from DNA oxidation during sequencing or FFPE-based DNA extraction using a filter-based method^69^. Subsequently, Strelka (RRID:SCR_005109)^70^ was used to identify small insertions or deletions. Finally, Oncotator (RRID:SCR_005183)^71^ was used to annotate the identified alterations.

### Ploidy and ploidy estimation

Tumor purity and ploidy were estimated using two orthogonal approaches. ABSOLUTE (RRID: SCR_005198)^72^ was applied for WES data, integrating variant allele frequency distributions together with segmented copy number profiles to infer absolute tumor purity, ploidy and the cancer cell fraction (CCF) of each somatic mutation, defines as the proportion of tumor cells harboring that mutation). Allelic segments corrected for tumor purity and ploidy were then used to derive allelic copy number. FACETS (RRID: SCR_026264)^73^ was applied to the same data to obtain joint estimates of purity, ploidy and allele-specific copy number.

For comparison between primary mucosal and primary cutaneous melanoma samples, we used FACETS estimates for both cohorts. Publicly available cutaneous melanoma data (TCGA-SKCM) were processed with FACETS and using one algorithm across both cohorts ensure that differences between them were not driven by the different algorithms.

For analysis within the mucosal melanoma cohort, we used ABSOLUTE estimates. Copy number prolife for ABSOLUTE were generated using panel of normals for coverage normalization and artefact filtering, and ABSOLUTE reports multiple candidate purity and ploidy solutions per sample, which were reviewed manually to select the most plausible call for each tumor. Selected solutions are reported in **Supplementary Table 2**.

### Heterogeneity and aneuploidy estimation

Intratumoral heterogeneity was quantified as the proportion of somatic mutations inferred to be subclonal within each tumor. Clonal mutations were defined as those with a CCF ≥ 0.8, whereas mutations with CCF < 0.8 were classified as subclonal, consistent with previous studies^50,74^. Aneuploidy was quantified as the fraction of the genome exhibiting allelic copy-number gain or loss based on the allelic segmentation described above.

### Mutational signatures calculation

De novo mutational signatures were generated in this cohort using an adaptation of non-negative matrix factorization^75^ via the Brunet update method^76^, as previously described in detail^77^, with the R package SomaticSignatures (RRID:SCR_025620)^67^ and non-negative matrix factorization (RRID:SCR_023124)^78^. Cosine similarity was used to compare the discovered signatures to the 30 existing discovered and validated signatures in COSMIC (RRID:SCR_002260)^79,80^, with a threshold of 0.85, and we also manually visualized and inspected similarities in mutational motifs between our signatures and COSMIC signatures.

### Copy number signature calculation

Copy number signature were derived from allele-specific segment profiles generated by FACETS. For each sample, segment coordinates together with major and minor allele copy number were extracted and combined into a single input profile. Segments were classified into the 4-channel copy number classification (CN48), which categorizes each segment by total copy number, heterozygosity status and segment length, using SigProfilerMatrixGenerator (v1.3.6; RRID: SCR_023122) as described previously^81^. The resulting CN48 matrix was fitted to the reference COSMIC v3.4 copy number signatures (GRCh37) using SigProfiler Assignment (v1.1.3; RRID:SCR_026899)^82^. Per-sample activities were normalized to the total number of assigned segments to yield the relative contribution of each signature, and signatures with zero contribution across all samples were excluded from downstream comparison.

### MM targeted sequencing data

Tumor genotyping of DFCI MM patients (n = 86) was performed by targeted sequencing of up to 447 genes (Oncopanel v15.0) of 86 MM patients that were previously published^22^. We focused on 23 gene alterations previously found to be enriched in MM^20^. For validation, AACR Project GENIE provided us with 420 MM patients from 14 medical centers. Of these, 110 patients lacking information for the alterations in 23 driver genes and 57 DFCI patients overlapping our discovery cohort were excluded, resulting in a nonoverlapping cohort (n=253 patients) (**Fig. 2a**)^83^.

### Targeted sequencing co-alteration analysis

The following variant types were assessed and included if present: HA (high amplification), 2DEL (loss of both copies), missense, splice, structural variation, nonsense, promoter mutation, truncation, Indel = Insertion and deletions, and Struct Var = Structural variations. The mutation and copy number alteration data for driver genes with CNA and mutational analysis (n = 420) was downloaded from cBioPortal^40^ on 06/04/2025. 110 samples that lacked profiling for one or more of the 23 selected driver genes were excluded, ensuring consistency in genetic profiles for analysis. The 51 samples from DFCI were also excluded to reduce the bias caused by redundant samples. Duplicate or serial samples from the same individual were excluded.

### Permutation analysis

Permuted datasets were generated using nonsynonymous mutations of 53 genes that were found in at least 5 MM patients from the GENIE cohort (N=203)^40^. To generate a null distribution, we permuted the dataset 1,000 times while maintaining the total number of mutations per donor and per gene, as well as arm-level somatic copy number status. We detected arm-level events via ASCETs software with default settings^84^. We determined the statistical significance of co-mutated gene pairs by comparing the observed number of co-mutated donors to the null distribution. To visualize significantly altered genes, we selected those altered in >10% of patients with a two-sided *P* value < 0.08, then performed hierarchical clustering using two-sided *P* values for gene combinations.

### Identification of genetic subtypes

R was used for all genetic subtype statistical analysis unless otherwise specified [version:4.3.3]. Python package CoMut^85^ was used to visualize the distribution of genetic alterations in patients. For each gene pair, ORs and Fisher’s exact *P* values were calculated in each cohort and in the combined cohort, with ORs derived from 2 × 2 contingency tables after adding a pseudo count of 0.5 to each cell (Haldane–Anscombe correction). Concordance between cohorts was assessed by plotting cohort-specific log2-transformed ORs. Cross-cohort significance was summarized using the product of the two cohort-specific P values, and gene pairs meeting the predefined significance threshold were highlighted in red. To visualize cohort-specific patterns, we also generated separate heat maps for each cohort, restricted to genes altered in >5% of patients and to gene pairs with OR > 3 or OR < 0.25. Because the distribution of anatomic sites of origin, stage, and genetic changes was remarkably similar between the two cohorts, the data were merged to provide a consolidated view, and statistical analyses were performed on the combined dataset. Principal component analysis (PCA) was performed to visualize the distribution of patients on the basis of genetic alterations. Pairwise associations among MM subtypes were assessed using ORs and Fisher’s exact test, with a *P* value less than 0.05 considered significant, unless otherwise explicitly stated. A ternary plot was used to visualize the occurrence of genetic subgroups in various tumor sites.

### Analysis of Clinical Characteristics by Genetic Subtype

Clinical data (demographics, molecular subtype, stage, recurrence pattern, treatment history, survival outcomes, selected comorbidities) were collected for 97 MM patients at DFCI to assess gene-clinical associations. Of these, 86 overlapped with the discovery cohort; For the additional 11 patients, they were classified as above by subtype. Sex distribution was compared between our mucosal melanoma cohort and a cutaneous melanoma cohort from cBioPortal (accessed 03/14/2026). Age at diagnosis was evaluated across mucosal melanoma genetic subtypes. The type of first recurrence compared across molecular subtypes. Stage at diagnosis, history of prior cancers, depression and anxiety, and brain metastasis were assessed for each subtype. To evaluate the effect of adjuvant therapy after surgery, overall recurrence-free survival (RFS) was calculated as the number of days between date of diagnosis and date of first recurrence for each subtype. Treatment categories used during the disease course were documented and grouped into four categories based on the immune checkpoint inhibitor backbone received: CTLA4 backbone, PD1/PD-L1 backbone, combined CTLA4/PD1/PD-L1 backbone, and Others (patients who did not receive any of these regimens). Subtype association with these treatment categories was then examined. Clinical response was also calculated for the first treatment in the metastatic setting.

### Bulk transcriptomic analysis

Bulk RNA-seq analysis were deconvolute using BayesPrism (RRID: SCR_027499)^86^, a Bayesian based method that leverages scRNA-seq reference profiles to estimate cellular composition and reconstruct cell-type specific gene expression from bulk RNAseq data. For mucosal melanoma. the single-cell transcriptomic reference was derived from our previously published mucosal melanoma scRNA-seq dataset^22^. For CM. the same deconvolution procedure was performed using melanoma scRNA-seq reference profiles from Tirosh et al^29^. Following deconvolution, estimated cell-type fractions and reconstructed tumor-cell-specific expression profiles were extracted for each sample. Tumor-specific expression profiles were used for downstream transcriptomic comparisons, with ana analyses between mucosal and CM restricted to genes shared across both datasets.

### Bulk RNAseq gene signature analysis

Gene signature analysis was performed using BayesPrism-reconstructed tumor cell expression profiles described above. Single-sample gene set enrichment analysis (ssGSEA) scores were calculated using GSVA R package (RRID:SCR_021058), and resulting gene signature scores were used for downstream comparative analyses. Gene sets included the MSigDB Hallmark^87^, TME functional gene signatures^88^ and previously defined melanoma tumor-state signatures^30,89,90^.

### Tissue imaging (Orion)

Tissue imaging with the RareCyte Orion platform (RRID:SCR_026855) was performed for all the specimens with the following protocol previously described by Lin et al^41^. In brief, tissue sections were deparaffinized and subjected to heat-induced epitope retrieval. Slides were incubated at 95 °C for 5 min and then at 105 °C for 5 min in EZ-AR2 Elegance buffer (BioGenex HX032). Autofluorescence was reduced by treatment with a peroxide-based quenching solution (4.5% H**₂**O**₂**, 25 mM NaOH in PBS) under illumination, followed by washes in surfactant-containing buffers. Slides were then incubated in Orion Signal Enhancer for 15 min, washed, and stained overnight at 4 °C with the multiplex antibody panel. After primary staining, sections were washed and incubated for 30 min with anti-DIG antibody and streptavidin to detect DIG- and biotin-labeled probes. Slides were washed and coverslipped with Orion mounting medium (RareCyte WA). For sequential H&E staining, Orion-stained sections were decoverslipped in PBS for overnight, followed by the standard H&E protocol.

### Orion image pre-processing and quality control

Assembly of raw CyCIF and Orion imaging data into a high-dimensional images, as well as nuclear segmentation and single-cell feature extraction, were performed using the open-source MCMICRO^91^ processing pipeline (RRID:SCR_022832), an open-source multiple-choice microscopy pipeline (version:38182748aa0ec021f684ce47248c57340d2f4cc7; full codes available at https://github.com/labsyspharm/mcmicro). The specific parameters used were optimized after iterative inspection of results, specifically focusing on performance of the segmentation module to ensure accurate identification of single cells (https://github.com/davidliu-lab/mucosal_melanoma_atlas_manuscript_code_2026). The quality of the segmentation was assessed, and the segmentation parameters were iteratively modified to improve the accuracy of the segmentation masks. After generating the segmentation masks, the mean fluorescence intensities of each marker for each cell were computed, resulting in a single-cell data table for each acquired whole-slide Orion image. The X/Y coordinates of annotated histologic regions on the whole slide image were used to extract the single-cell data of cells that lie within the annotated regions. Multiple approaches were also taken to ensure the quality of the single-cell data. At the image level, the cross-cycle image registration and tissue integrity were reviewed; regions that were poorly registered or contained severely deformed tissues and artifacts were identified, and cells inside those regions were excluded. Antibody staining patterns that gave low confidence by visual evaluation were also excluded from the analyses. Panels, abbreviations, and key properties of these markers are described in **Supplementary Table 16**.

### Orion single-cell gating-based phenotyping

Cells from Orion data were phenotyped using a gating-based classification approach as previously described.^92^ In brief, gates for each marker were defined using an open-source visualization tool (https://github.com/labsyspharm/gater). These gates were then used to rescale single-cell expression values between 0 and 1, with values above 0.5 indicating marker positivity. The normalized data were subsequently used for cell-type calling and unsupervised clustering. Phenotype labels were assigned using the SCIMAP^93^ Python package (RRID:SCR_024751) based on a hierarchical classification of marker expression patterns (**Extended Data Fig. 1b**). Assigned cell types were validated by overlaying phenotype labels onto the corresponding images.

### Tumor boundary definition and quantification of tumor-infiltrating immune cell proportions

Tumor boundaries were defined using the SpatialCells package (https://github.com/SemenovLab/SpatialCells)^94^. Single-cell centroid coordinates of SOX10⁺ tumor cells were used to delineate tumor boundaries using density-based spatial clustering (DBSCAN), followed by polygon-based boundary reconstruction to define contiguous tumor regions. After tumor boundaries were defined, the proportion of each cell type within the tumor was calculated by normalizing the number of cells of that cell type to the total number of cells within the defined tumor region. For tumors lacking SOX10 signal on the Orion image but with tumor identified by a pathologist on the post-Orion H&E or adjacent tissue section, the pathologist-defined tumor region was used to delineate the corresponding tumor region on the ORION image.

### Orion neighborhood spatial analysis

Cell centroid coordinates were converted to microns. A Delaunay triangulation was performed on the coordinates of the target cell population, and edges were adaptively filtered based on local nearest-neighbor distance to retain short-range connections. Cells located within 50 μm of any target cell were defined as neighbors.

### scRNA-seq analysis

scRNA-seq data were obtained from our previously published mucosal melanoma cohort^22^ (GEO: GSE270464 and GSE269936), in which alignment, ambient RNA correction, doublet removal, quality filtering and cell-type annotation were performed as described. Scanpy (v1.11.2; RRID: SCR_0181139)^95^ was used for all subsequent processing and analysis. Cells with fewer than 200 detected genes or more than 20% of reads mapping to mitochondrial genes were removed, and genes detected in fewer than three cells were excluded. Analyses were restricted to eight patients with progression sampled after immunotherapy. Counts were normalized 10,000 per cells and log-transformed, the 5,000 most highly variable genes were selected, and expression values were scaled to a maximum of 10 before principal component analysis (PCA). The first 30 principal components and 15 nearest neighbors were used for Uniform Manifold Approximation and Projection (UMAP) visualization and Leiden clustering.

The myeloid and lymphoid compartments annotated in the original study were extracted and reprocessed independently. For each compartment, raw counts were re-normalized and log-transformed, expression values were scaled without zero-centering, and PCA, neighborhood graph construction (30 principal components, 15 nearest neighbors), UMAP and Leiden clustering were performed as above at resolutions of 0.1-0.4. Myeloid clusters were annotated at resolution 0.2 as macrophages (*CD68*, *CD163*, *C1QA–C*, *APOE*, *MSR1*, *MERTK*), monocytes (*FCN1*, *VCAN*, *S100A8/9*, *LYZ*), conventional dendritic cells (*CD1C*, *CLEC10A*, *FCER1A*), plasmacytoid dendritic cells (*IL3RA*, *TCF4*, *IRF7*), neutrophils (*CSF3R*, *FCGR3B*) and mast cells (*KIT*, *TPSAB1*, *TPSB2*). Lymphoid clusters were annotated at resolution 0.3 as CD8⁺ T cells (*CD8A*, *CD8B*, *GZMB*), CD4⁺ T cells (*CD4*, *CTLA4*), regulatory T cells (*FOXP3*), natural killer cells (*KLRD1*, *NCAM1*) and B cells (*MS4A1*, *CD79A*, *CD19*); clusters expressing *MLANA* or *PMEL* were considered tumor-contaminated and excluded. Lymphoid clusters were annotated at resolution 0.3 as regulatory T cells (*FOXP3*, *CTLA4*, *TIGIT*, *IL2RA*), and non-regulatory T cells (*CD3D*, *CD3E*, *CD4*, *CD8A*, *CD8B*), natural killer (NK) cells (*KLRD1*, *NCAM1*) and B/plasma cells (*MS4A1*, *CD79A*, *CD19*), based on canonical marker expression and cluster-level gene-expression patterns; one cluster could not be confidently assigned and was labelled unknown. Clusters expressing *MLANA* or *PMEL* were assessed for tumor contamination.

### Macrophage polarization and transcriptional subsets

Macrophages were extracted from the myeloid compartment and reprocessed as above. Pro-inflammatory (M1-like) and immunosuppressive (M2-like) polarization scores were calculated per cell from published gene signatures (**Extended Data Table 14**)^74^ using sc.tl.score_genes(). Cells scoring below zero for both programs were classified as “other”. For the remaining cells, a polarization score were classified as the pro-inflammatory score minus the immunosuppressive score; cells above the 85^th^ percentile of this distribution were classified as the pro-inflammatory, those below the 15^th^ percentile as immunosuppressive, and the remainder as mixed. Macrophages were additionally scored for five TAM programs — inflammatory, interferon-responsive, *SPP1*/*APOE*, regulatory and resident tissue macrophage-like TAM^96^ — and scores were compared between M1-like and M2-like macrophages using two-sided Mann-Whitney *U* tests.

### Ligand-receptor interactions analysis

Cell–cell ligand–receptor interactions were inferred using CellPhoneDB (v2.1.7; RRID:SCR_017054)^97^, with the refined cell-type annotations and raw scRNA-seq counts as input. Statistical significance was assessed using the permutation-based framework implemented in CellPhoneDB (1000 permutations), and interactions were considered significant at *P* < 0.05. Analyses focused on interactions between macrophage subsets and lymphoid populations.

### Statistical tests

All statistical comparisons between groups were performed using the Mann-Whitney U rank test to assess differences in the distribution of values between groups, implemented via the mannwhitneyu function in the SciPy55 Python package (RRID: SCR_008058) or the stats and ggpubr R packages (RRID:SCR_021139). Paired comparisons were generally performed using two-sided Willcoxon signed-rank tests. For the paired primary to metastatic tumor infiltrated immune cell proportion comparison shown in **Fig. 3e** and **Extended Data Fig. 5e**, only five paired samples were available; because the minimum attainable exact two-sided *P* value for a Wilcoxon signed-rank test with five non-zero pairs is 0.0625, two-sided paired *t*-tests were used for these comparisons.

## REFERENCES

1. Yde, S. S., Sjoegren, P., Heje, M. & Stolle, L. B. Mucosal Melanoma: a Literature Review. Curr Oncol Rep 20, 28 (2018).

2. Bishop, K. D. & Olszewski, A. J. Epidemiology and survival outcomes of ocular and mucosal melanomas: A population-based analysis. Intl Journal of Cancer 134, 2961–2971 (2014).

3. Cui, C. et al. An Evidence-Based Staging System for Mucosal Melanoma: A Proposal. Ann Surg Oncol 29, 5221–5234 (2022).

4. Altieri, L., Eguchi, M., Peng, D. H. & Cockburn, M. Predictors of mucosal melanoma survival in a population-based setting. Journal of the American Academy of Dermatology 81, 136–142.e2 (2019).

5. Mucosal Melanoma: New Insights and Therapeutic Options for a Unique and Aggressive Disease | CancerNetwork.

6. Bachar, G. et al. Mucosal melanomas of the head and neck: The Princess Margaret Hospital experience. Head & Neck 30, 1325–1331 (2008).

7. Guo, J. et al. Phase II, Open-Label, Single-Arm Trial of Imatinib Mesylate in Patients With Metastatic Melanoma Harboring *c-Kit* Mutation or Amplification. JCO 29, 2904–2909 (2011).

8. Hodi, F. S. et al. Imatinib for Melanomas Harboring Mutationally Activated or Amplified *KIT* Arising on Mucosal, Acral, and Chronically Sun-Damaged Skin. JCO 31, 3182–3190 (2013).

9. Shi, C. et al. The efficacy and safety of dalpiciclib, a cyclin-dependent kinase 4/6 inhibitor, in patients with advanced head and neck mucosal melanoma harboring CDK4 amplification. BMC Med 22, 215 (2024).

10. Han, Y. et al. Repurposing Ponatinib as a Potent Agent against *KIT* Mutant Melanomas. Theranostics 9, 1952–1964 (2019).

11. D’Angelo, S. P. et al. Efficacy and Safety of Nivolumab Alone or in Combination With Ipilimumab in Patients With Mucosal Melanoma: A Pooled Analysis. JCO 35, 226–235 (2017).

12. Uhara, H. et al. Five-year survival with nivolumab in previously untreated Japanese patients with advanced or recurrent malignant melanoma. The Journal of Dermatology 48, 592–599 (2021).

13. Dimitriou, F. et al. Single-agent anti-PD-1 or combined with ipilimumab in patients with mucosal melanoma: an international, retrospective, cohort study. Annals of Oncology 33, 968–980 (2022).

14. Rose, A. A. N. et al. Biologic subtypes of melanoma predict survival benefit of combination anti-PD1+anti-CTLA4 immune checkpoint inhibitors versus anti-PD1 monotherapy. J Immunother Cancer 9, e001642 (2021).

15. Larkin, J. et al. Five-Year Survival with Combined Nivolumab and Ipilimumab in Advanced Melanoma. N Engl J Med 381, 1535–1546 (2019).

16. Hamid, O. et al. Antitumour activity of pembrolizumab in advanced mucosal melanoma: a post-hoc analysis of KEYNOTE-001, 002, 006. Br J Cancer 119, 670–674 (2018).

17. Hayward, N. K. et al. Whole-genome landscapes of major melanoma subtypes. Nature 545, 175–180 (2017).

18. Newell, F. et al. Comparative Genomics Provides Etiologic and Biological Insight into Melanoma Subtypes. Cancer Discovery 12, 2856–2879 (2022).

19. Nassar, K. W. & Tan, A. C. The mutational landscape of mucosal melanoma. Seminars in Cancer Biology 61, 139–148 (2020).

20. Newell, F. et al. Whole-genome landscape of mucosal melanoma reveals diverse drivers and therapeutic targets. Nat Commun 10, 3163 (2019).

21. Buchbinder, E. I. et al. Characterization of genetics in patients with mucosal melanoma treated with immune checkpoint blockade. Cancer Medicine 10, 2627–2635 (2021).

22. Babu, S. et al. Specific oncogene activation of the cell of origin in mucosal melanoma. Preprint at 10.1101/2024.04.22.590595 (2024).

23. Hintzsche, J. D. et al. Whole-exome sequencing identifies recurrent SF3B1 R625 mutation and comutation of NF1 and KIT in mucosal melanoma. Melanoma Research 27, 189–199 (2017).

24. Curtin, J. A., Patel, H. N., Cho, K.-H. & LeBoit, P. E. Distinct Sets of Genetic Alterations in Melanoma. n engl j med (2005).

25. Hayward, N. K. et al. Whole-genome landscapes of major melanoma subtypes. Nature 545, 175–180 (2017).

26. Hoadley, K. A. et al. Cell-of-Origin Patterns Dominate the Molecular Classification of 10,000 Tumors from 33 Types of Cancer. Cell 173, 291–304.e6 (2018).

27. Broit, N. et al. Meta-Analysis and Systematic Review of the Genomics of Mucosal Melanoma. Molecular Cancer Research 19, 991–1004 (2021).

28. Hoek, K. S., et al. *In vivo* Switching of Human Melanoma Cells between Proliferative and Invasive States. Cancer Research 68, 650–656 (2008).

29. Tirosh, I. et al. Dissecting the multicellular ecosystem of metastatic melanoma by single-cell RNA-seq. Science 352, 189–196 (2016).

30. Tsoi, J. et al. Multi-stage Differentiation Defines Melanoma Subtypes with Differential Vulnerability to Drug-Induced Iron-Dependent Oxidative Stress. Cancer Cell 33, 890–904.e5 (2018).

31. Andrews, M. C. et al. Multi-modal molecular programs regulate melanoma cell state. Nat Commun 13, 4000 (2022).

32. De Visser, K. E. & Joyce, J. A. The evolving tumor microenvironment: From cancer initiation to metastatic outgrowth. Cancer Cell 41, 374–403 (2023).

33. Giraldo, N. A. et al. The clinical role of the TME in solid cancer. Br J Cancer 120, 45–53 (2019).

34. Bejarano, L., Jordāo, M. J. C. & Joyce, J. A. Therapeutic Targeting of the Tumor Microenvironment. Cancer Discovery 11, 933–959 (2021).

35. Binnewies, M. et al. Understanding the tumor immune microenvironment (TIME) for effective therapy. Nat Med 24, 541–550 (2018).

36. Vos, J. L. et al. Diversity of the immune microenvironment and response to checkpoint inhibitor immunotherapy in mucosal melanoma. JCI Insight 9, e179982 (2024).

37. Nakamura, Y. et al. Poor Lymphocyte Infiltration to Primary Tumors in Acral Lentiginous Melanoma and Mucosal Melanoma Compared to Cutaneous Melanoma. Front. Oncol. 10, 524700 (2020).

38. Shteinman, E. R. et al. Enhanced measures of neoantigenicity capture unique tumor-immune interactions across primary melanoma subtypes. Genome Med 18, 106 (2026).

39. Li, Y. et al. Single-Cell Transcriptomic Landscape Deciphers Intratumoral Heterogeneity and Subtypes of Acral and Mucosal Melanomas. Clinical Cancer Research 31, 2495–2514 (2025).

40. The AACR Project GENIE Consortium et al. AACR Project GENIE: Powering Precision Medicine through an International Consortium. Cancer Discovery 7, 818–831 (2017).

41. Lin, J.-R. et al. High-plex immunofluorescence imaging and traditional histology of the same tissue section for discovering image-based biomarkers. Nat Cancer 4, 1036–1052 (2023).

42. Akbani, R. et al. Genomic Classification of Cutaneous Melanoma. Cell 161, 1681–1696 (2015).

43. Gulve, N. et al. DAXX-ATRX regulation of p53 chromatin binding and DNA damage response. Nat Commun 13, 5033 (2022).

44. Oppel, F. et al. Loss of atrx cooperates with p53-deficiency to promote the development of sarcomas and other malignancies. PLoS Genet 15, e1008039 (2019).

45. Lehrer, S., Rheinstein, P. H., Green, S. & Rosenzweig, K. E. von Willebrand Factor Gene Expression in Primary Lower Grade Glioma: Mutually Co-Occurring Mutations in von Willebrand Factor, ATRX, and TP53. Brain Tumor Res Treat 7, 33 (2019).

46. Malumbres, M. & Barbacid, M. Cell cycle, CDKs and cancer: a changing paradigm. Nat Rev Cancer 9, 153–166 (2009).

47. Quek, C. et al. mucosal melanoma identified in a study of 27 Australian mucosal.

48. Nagarajan, P. et al. Prognostic model for patient survival in primary anorectal mucosal melanoma: stage at presentation determines relevance of histopathologic features. Modern Pathology 33, 496–513 (2020).

49. Altieri, L., Wong, M. K., Peng, D. H. & Cockburn, M. Mucosal melanomas in the racially diverse population of California. Journal of the American Academy of Dermatology 76, 250– 257 (2017).

50. Tarantino, G. et al. Genomic heterogeneity and ploidy identify patients with intrinsic resistance to PD-1 blockade in metastatic melanoma. ScieNce AdvANceS (2024).

51. Zhou, R. et al. Analysis of Mucosal Melanoma Whole-Genome Landscapes Reveals Clinically Relevant Genomic Aberrations. Clinical Cancer Research 25, 3548–3560 (2019).

52. Kim, D. W. et al. Clinicopathological features and clinical outcomes associated with *TP53* and *BRAF* ^*N on-V* 600^ mutations in cutaneous melanoma patients. Cancer 123, 1372–1381 (2017).

53. Said, R. et al. Characteristics and survival of patients with advanced cancer and p53 mutations. Oncotarget 5, 3871–3879 (2014).

54. Vos, J. L. et al. Diversity of the immune microenvironment and response to checkpoint inhibitor immunotherapy in mucosal melanoma. JCI Insight 9, e179982 (2024).

55. Chohan, M. H., Perry, M., Laurance-Young, P., Salih, V. M. & Foey, A. D. Prognostic Role of CD68+ and CD163+ Tumour-Associated Macrophages and PD-L1 Expression in Oral Squamous Cell Carcinoma: A Meta-Analysis. Br J Biomed Sci 80, 11065 (2023).

56. Yang, Y., Sun, D., Ma, X., Wang, T. & Wu, J. CD68- and CD163-positive tumor-associated macrophages in renal clear cell carcinoma. Sci Rep 15, 35138 (2025).

57. Wang, C. et al. The Prognostic and Clinical Value of Tumor-Associated Macrophages in Patients With Breast Cancer: A Systematic Review and Meta-Analysis. Front. Oncol. 12, 905846 (2022).

58. Kinoshita, J. et al. Prognostic value of tumor-infiltrating CD163+macrophage in patients with metastatic gastric cancer undergoing multidisciplinary treatment. BMC Cancer 22, 608 (2022).

59. Cheng, L.-J. et al. Effect of the TIM-3/Gal-9 signaling pathway on macrophage polarization in peri-implantitis. PLoS One 20, e0328258 (2025).

60. Kandel, S., Adhikary, P., Li, G. & Cheng, K. The TIM3/Gal9 signaling pathway: An emerging target for cancer immunotherapy. Cancer Letters 510, 67–78 (2021).

61. Díaz-García, E. et al. Immune Checkpoint Biomarkers Galectin-9 and TIM-3 Predict Melanoma and Lung Cancer Mortality in Obstructive Sleep Apnoea. Archivos de Bronconeumología 61, 735–748 (2025).

62. Yang, R. et al. Galectin-9 interacts with PD-1 and TIM-3 to regulate T cell death and is a target for cancer immunotherapy. Nat Commun 12, 832 (2021).

63. Mehta, A. et al. Immunotherapy Resistance by Inflammation-Induced Dedifferentiation. Cancer Discovery 8, 935–943 (2018).

64. Landsberg, J. et al. Melanomas resist T-cell therapy through inflammation-induced reversible dedifferentiation. Nature 490, 412–416 (2012).

65. Hossain, S. M. et al. Innate immune checkpoint inhibitor resistance is associated with melanoma sub-types exhibiting invasive and de-differentiated gene expression signatures. Front. Immunol. 13, 955063 (2022).

66. Sade-Feldman, M. et al. Resistance to checkpoint blockade therapy through inactivation of antigen presentation. Nat Commun 8, 1136 (2017).

67. Gehring, J. S., Fischer, B., Lawrence, M. & Huber, W. SomaticSignatures: inferring mutational signatures from single-nucleotide variants. Bioinformatics 31, 3673–3675 (2015).

68. Cibulskis, K. et al. Sensitive detection of somatic point mutations in impure and heterogeneous cancer samples. Nat Biotechnol 31, 213–219 (2013).

69. Costello, M. et al. Discovery and characterization of artifactual mutations in deep coverage targeted capture sequencing data due to oxidative DNA damage during sample preparation. Nucleic Acids Research 41, e67–e67 (2013).

70. Saunders, C. T. et al. Strelka: accurate somatic small-variant calling from sequenced tumor–normal sample pairs. Bioinformatics 28, 1811–1817 (2012).

71. Ramos, A. H. et al. Oncotator: Cancer Variant Annotation Tool. Human Mutation 36, E2423–E2429 (2015).

72. Carter, S. L. et al. Absolute quantification of somatic DNA alterations in human cancer. Nat Biotechnol 30, 413–421 (2012).

73. Shen, R. & Seshan, V. E. FACETS: allele-specific copy number and clonal heterogeneity analysis tool for high-throughput DNA sequencing. Nucleic Acids Res 44, e131 (2016).

74. Liu, H. et al. Integrative molecular and spatial analysis reveals evolutionary dynamics and tumor-immune interplay of in situ and invasive acral melanoma. Cancer Cell 42, 1067–1085.e11 (2024).

75. Lee, D. D. & Seung, H. S. Learning the parts of objects by non-negative matrix factorization. Nature 401, 788–791 (1999).

76. Brunet, J.-P., Tamayo, P., Golub, T. R. & Mesirov, J. P. Metagenes and molecular pattern discovery using matrix factorization. Proc. Natl. Acad. Sci. U.S.A. 101, 4164–4169 (2004).

77. Liu, D. et al. Mutational patterns in chemotherapy resistant muscle-invasive bladder cancer. Nat Commun 8, 2193 (2017).

78. Gaujoux, R. & Seoighe, C. A flexible R package for nonnegative matrix factorization. BMC Bioinformatics 11, 367 (2010).

79. Forbes, S. A. et al. COSMIC: exploring the world’s knowledge of somatic mutations in human cancer. Nucleic Acids Research 43, D805–D811 (2015).

80. Australian Pancreatic Cancer Genome Initiative et al. Signatures of mutational processes in human cancer. Nature 500, 415–421 (2013).

81. Bergstrom, E. N. et al. SigProfilerMatrixGenerator: a tool for visualizing and exploring patterns of small mutational events. BMC Genomics 20, 685 (2019).

82. Díaz-Gay, M. et al. 1 Assigning mutational signatures to individual samples and individual somatic 2 mutations with SigProfilerAssignment.

83. Cerami, E. et al. The cBio Cancer Genomics Portal: An Open Platform for Exploring Multidimensional Cancer Genomics Data. Cancer Discovery 2, 401–404 (2012).

84. Spurr, L. F. et al. Quantification of aneuploidy in targeted sequencing data using ASCETS. Bioinformatics 37, 2461–2463 (2021).

85. Crowdis, J., He, M. X., Reardon, B. & Van Allen, E. M. CoMut: visualizing integrated molecular information with comutation plots. Bioinformatics 36, 4348–4349 (2020).

86. Chu, T., Wang, Z., Pe’er, D. & Danko, C. G. Cell type and gene expression deconvolution with BayesPrism enables Bayesian integrative analysis across bulk and single-cell RNA sequencing in oncology. Nat Cancer 3, 505–517 (2022).

87. Subramanian, A., et al. Gene set enrichment analysis: A knowledge-based approach for interpreting genome-wide expression profiles. Proc. Natl. Acad. Sci. U.S.A. 102, 15545– 15550 (2005).

88. Bagaev, A. et al. Conserved pan-cancer microenvironment subtypes predict response to immunotherapy. Cancer Cell 39, 845–865.e7 (2021).

89. Wouters, J. et al. Robust gene expression programs underlie recurrent cell states and phenotype switching in melanoma. Nat Cell Biol 22, 986–998 (2020).

90. Pozniak, J. et al. A TCF4-dependent gene regulatory network confers resistance to immunotherapy in melanoma. Cell 187, 166–183.e25 (2024).

91. Schapiro, D. et al. MCMICRO: a scalable, modular image-processing pipeline for multiplexed tissue imaging. Nat Methods 19, 311–315 (2022).

92. Shi, Y. Immunogenic tumor mass dormancy as a driver of persistent residual lesions in immunotherapy-treated melanoma.

93. Nirmal, A. J. & Sorger, P. K. SCIMAP: A Python Toolkit for Integrated SpatialAnalysis of Multiplexed Imaging Data. JOSS 9, 6604 (2024).

94. Wan, G. et al. SpatialCells: automated profiling of tumor microenvironments with spatially resolved multiplexed single-cell data. Briefings in Bioinformatics 25, bbae189 (2024).

95. Wolf, F. A., Angerer, P. & Theis, F. J. SCANPY: large-scale single-cell gene expression data analysis. Genome Biol 19, 15 (2018).

96. Ma, R.-Y., Black, A. & Qian, B.-Z. Macrophage diversity in cancer revisited in the era of single-cell omics. Trends in Immunology 43, 546–563 (2022).

97. Vento-Tormo, R. et al. Single-cell reconstruction of the early maternal–fetal interface in humans. Nature 563, 347–353 (2018).

