## Supplementary Figures for "Genomic classification and immune profiling define mucosal melanoma"

**EXTENDED FIGURES:**

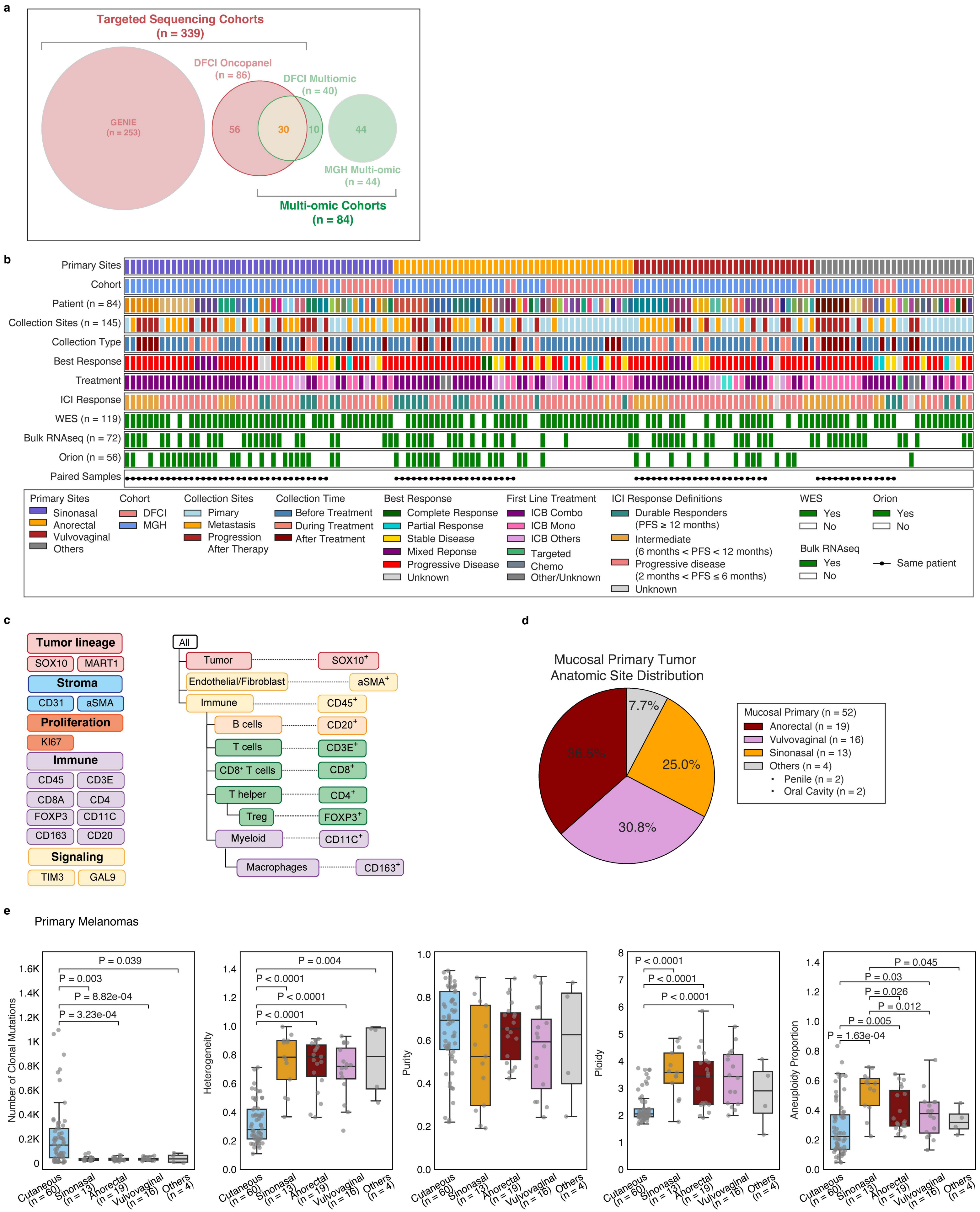
a
Targeted Sequencing Cohorts
(n = 339)
DFCI Oncopanel
(n = 86)
DFCI Multiomic
(n = 40)
GENIE
(n = 253)
56
30
10
44
MGH Multi-omic
(n = 44)
Multi-omic Cohorts
(n = 84)
b
Primary Sites
Cohort
Patient (n = 84)
Collection Sites (n = 145)
Collection Type
Best Response
Treatment
ICI Response
WES (n = 119)
Bulk RNAseq (n = 72)
Orion (n = 56)
Paired Samples
Primary Sites
Sinonasal
Anorectal
Vulvovaginal
Others
Cohort
DFCI
MGH
Collection Sites
Pimary
Metastasis
Progression After Therapy
Collection Time
Before Treatment
During Treatment
After Treatment
Best Response
Complete Response
Partial Response
Stable Disease
Mixed Reponse
Progressive Disease
Unknown
First Line Treatment
ICB Combo
ICB Mono
ICB Others
Targeted
Chemo
Other/Unknown
ICI Response Definitions
Durable Responders (PFS ≥ 12 months)
Intermediate (6 months < PFS < 12 months)
Progressive disease (2 months < PFS ≤ 6 months)
Unknown
WES
Yes
No
Bulk RNAseq
Yes
No
Orion
Yes
No
Same patient
c
Tumor lineage
SOX10
MART1
Stroma
CD31
aSMA
Proliferation
KI67
Immune
CD45
CD3E
CD8A
CD4
FOXP3
CD11C
CD163
CD20
Signaling
TIM3
GAL9
All
Tumor
SOX10+
Endothelial/Fibroblast
aSMA+
Immune
CD45+
B cells
CD20+
T cells
CD3E+
CD8+ T cells
CD8+
T helper
CD4+
Treg
FOXP3+
Myeloid
CD11C+
Macrophages
CD163+
d
Mucosal Primary Tumor
Anatomic Site Distribution
7.7%
36.5%
25.0%
30.8%
Mucosal Primary (n = 52)
Anorectal (n = 19)
Vulvovaginal (n = 16)
Sinonasal (n = 13)
Others (n = 4)
• Penile (n = 2)
• Oral Cavity (n = 2)
e
Primary Melanomas
Number of Clonal Mutations
P = 0.039
P = 0.003
P = 8.82e-04
P = 3.23e-04
Heterogeneity
P = 0.004
P < 0.0001
P < 0.0001
P < 0.0001
Purity
Ploidy
P < 0.0001
P < 0.0001
P < 0.0001
Aneuploidy Proportion
P = 0.045
P = 0.03
P = 0.026
P = 0.012
P = 0.005
P = 1.63e-04
Cutaneous (n = 60)
Sinonasal (n = 13)
Anorectal (n = 19)
Vulvovaginal (n = 16)
Others (n = 4)

**Extended Data Fig. 1:** **a**, Composition of patient cohorts analyzed in this study. Targeted sequencing cohorts (*n* = 339 patients) comprise the AACR project GENIE cohort (*n* = 253) and the DFCI OncoPanel cohort (*n* = 86). Multi-omic cohorts (*n* = 84 patients) comprise the DFCI (*n* = 40) and MGH (*n* = 44) cohorts. Thirty patients profiled by OncoPanel were also included in the DFCI multi-omic cohort, giving 393 unique patients in total. Circle area is proportional to cohort size; overlap indicates shared patients. **b**, Overview of the multi-omic MM cohort. Each column represents one tumor sample (*n* = 145 samples from *n* = 84 patients). Tracks indicate contributing institution (Cohort), patient identity, anatomic context (SITE), timing of collection relative to therapy (COLLECTION), best overall response (BOR), and therapy class (CATEGORY). The lower three tracks indicate availability of whole-exome sequencing (WES), bulk RNA sequencing, and multiplexed imaging (Orion) for each sample. **c**, Antibody panel and cell-type classification hierarchy used for multiplexed imaging. Left, markers grouped by biological category. Right, the gating hierarchy by which single cells were assigned to lineage and subset identities; superscript symbols denote marker positivity. **d**, Distribution of anatomic sites of origin among primary mucosal melanoma tumors. "Others" comprises penile (*n* = 2) and oral cavity (*n* = 2) cases. **e**, Comparison of genomic features between primary CM and primary MM stratified by anatomic site of origin. Box plots show the median and interquartile range (IQR), with whiskers extending to 1.5 × IQR and individual tumors shown as points. Each point represents one tumor. Groups were compared using two-sided Mann-Whitney *U* tests; only significant comparisons are annotated ($P < 0.05$).

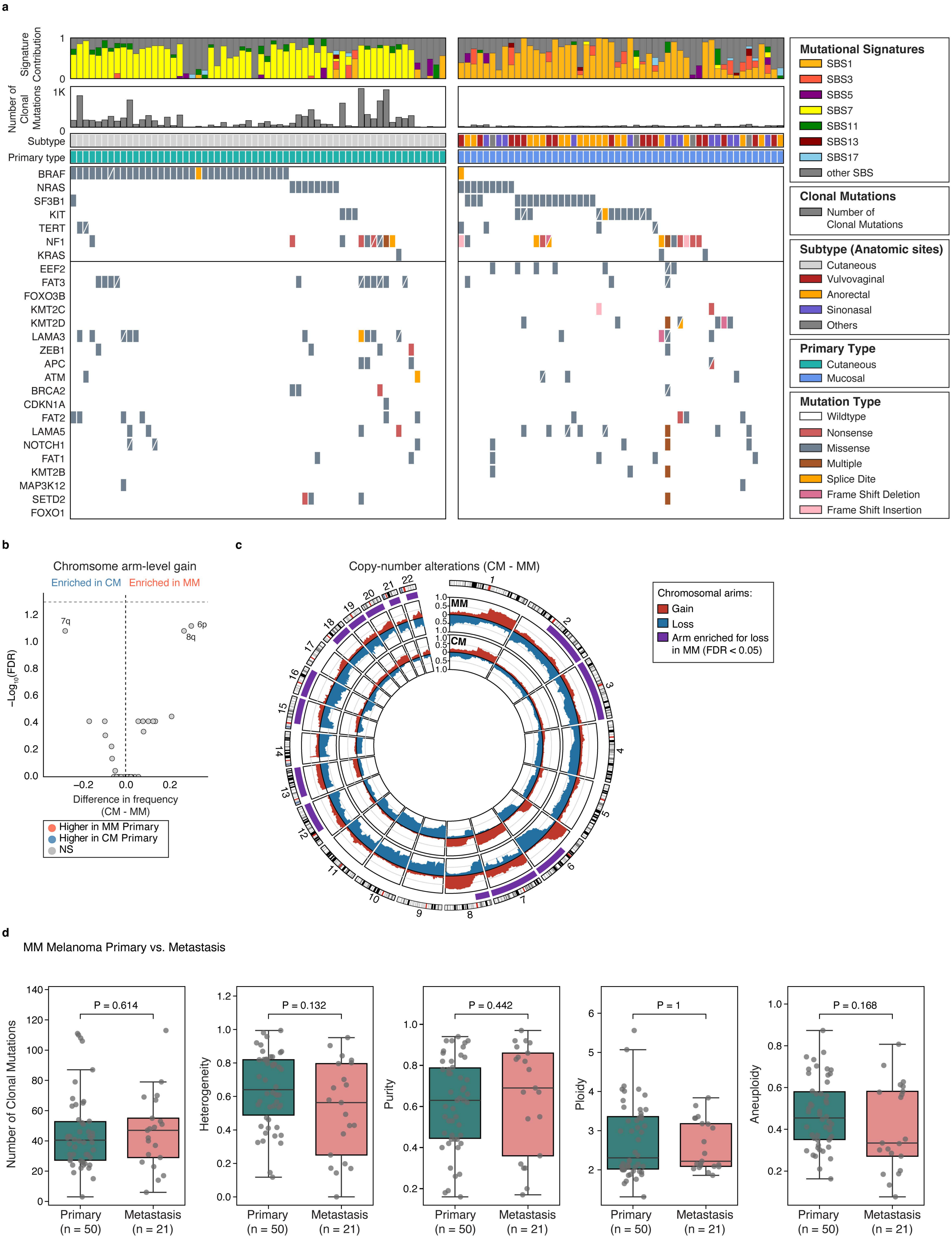

a
Signature Contribution
Number of Clonal Mutations
1K
Subtype
Primary type
BRAF
NRAS
SF3B1
KIT
TERT
NF1
KRAS
EEF2
FAT3
FOXO3B
KMT2C
KMT2D
LAMA3
ZEB1
APC
ATM
BRCA2
CDKN1A
FAT2
LAMA5
NOTCH1
FAT1
KMT2B
MAP3K12
SETD2
FOXO1
Mutational Signatures
SBS1
SBS3
SBS5
SBS7
SBS11
SBS13
SBS17
other SBS
Clonal Mutations
Number of Clonal Mutations
Subtype (Anatomic sites)
Cutaneous
Vulvovaginal
Anorectal
Sinonasal
Others
Primary Type
Cutaneous
Mucosal
Mutation Type
Wildtype
Nonsense
Missense
Multiple
Splice Dite
Frame Shift Deletion
Frame Shift Insertion
b
Chromsome arm-level gain
Enriched in CM
Enriched in MM
7q
6p
8q
−Log10(FDR)
Difference in frequency (CM - MM)
Higher in MM Primary
Higher in CM Primary
NS
c
Copy-number alterations (CM - MM)
MM
CM
Chromosomal arims:
Gain
Loss
Arm enriched for loss in MM (FDR < 0.05)
d
MM Melanoma Primary vs. Metastasis
P = 0.614
Number of Clonal Mutations
P = 0.132
Heterogeneity
P = 0.442
Purity
P = 1
Ploidy
P = 0.168
Aneuploidy
Primary (n = 50)
Metastasis (n = 21)

**Extended Data Fig. 2:** **a**, Genomic landscape of primary MM (left) and primary MM (right) melanoma. Each column represents one tumor. Top, relative contribution of SBS signatures per tumor, Signatures 1, 3, 5, 7, 11, 13 and 17 are shown individually; all remaining signatures and the unassigned residual are grouped as “other SBS”. Second, number of clonal mutations per tumor. Third and fourth tracks indicate anatomic site of origin and disease subtype. Lower panel, per-tumor alterations in melanoma driver genes (upper block) and additional recurrently altered genes (lower block), colored by mutation type. Genes are ordered by alteration frequency within each block. **b**, Comparison of chromosome arm-level copy-number gain between primary CM and MM. The *x* axis shows the difference in the proportion of tumors with arm-level gain (primary CM – primary MM) and the *y* axis shows the FDR-adjust *P* value. *P* values were calculated using two-sided Fisher’s exact tests with Benjamini-Hochberg correction; the dashed line indicates an FDR-adjusted *P* value of 0.05. **c**, Genome-wide copy number alteration frequency in primary MM (outer track) and primary CM (inner track). For each 1-Mb bin, bars show the proportion of tumors with a gain (red, outward) or a loss (blue, inward). Purple arcs indicate chromosome arms significantly enriched for loss in primary MM (FDR-adjusted $P < 0.05$; Fig. 1e). **d**, Genomic features comparison between primary ($n = 50$) and metastasis ($n = 21$) MM tumors. Shown are number of clonal mutations, intratumoral heterogeneity, purity, ploidy and aneuploidy derived from WES. Box plots show the median and interquartile range (IQR), with whiskers extending to 1.5 ´ IQR and individual tumors shown as points. Groups were compared using two-sided Mann-Whitney *U* tests; significant *P* values ($P < 0.05$) are shown.

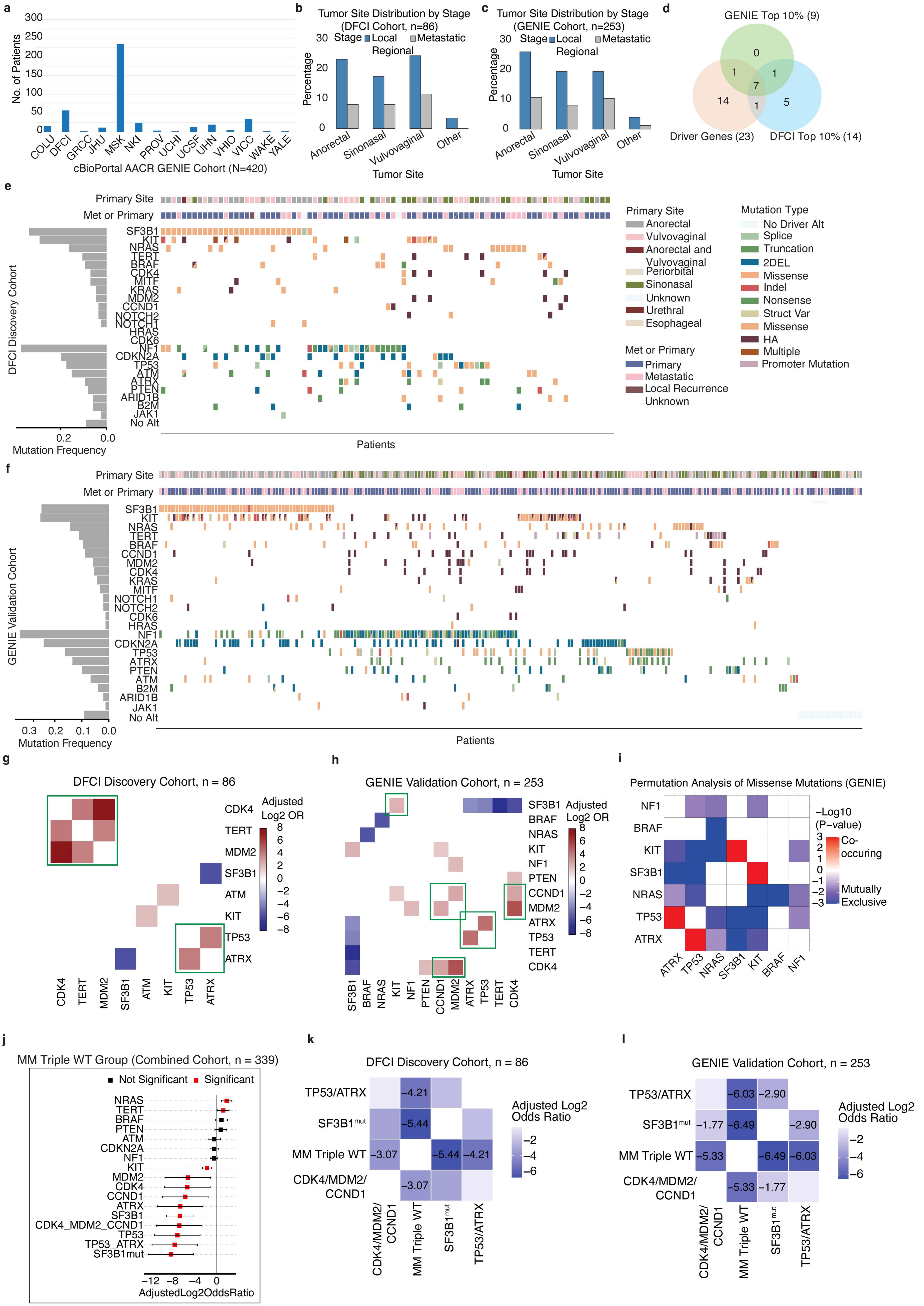
a
No. of Patients
COLU
DFCI
GRCC
JHU
MSK
NKI
PROV
UCHI
UCSF
UHN
VHIO
VICC
WAKE
YALE
cBioPortal AACR GENIE Cohort (N=420)
b
Tumor Site Distribution by Stage
(DFCI Cohort, n=86)
Stage
Local
Regional
Metastatic
Percentage
Anorectal
Sinonasal
Vulvovaginal
Other
Tumor Site
c
Tumor Site Distribution by Stage
(GENIE Cohort, n=253)
d
GENIE Top 10% (9)
Driver Genes (23)
DFCI Top 10% (14)
e
Primary Site
Met or Primary
DFCI Discovery Cohort
Mutation Frequency
Patients
Primary Site
Anorectal
Vulvovaginal
Anorectal and Vulvovaginal
Periorbital
Sinonasal
Unknown
Urethral
Esophageal
Met or Primary
Primary
Metastatic
Local Recurrence
Unknown
Mutation Type
No Driver Alt
Splice
Truncation
2DEL
Missense
Indel
Nonsense
Struct Var
Missense
HA
Multiple
Promoter Mutation
f
GENIE Validation Cohort
No Alt
g
DFCI Discovery Cohort, n = 86
Adjusted Log2 OR
h
GENIE Validation Cohort, n = 253
i
Permutation Analysis of Missense Mutations (GENIE)
−Log10 (P−value)
Co-occuring
Mutually Exclusive
j
MM Triple WT Group (Combined Cohort, n = 339)
Not Significant
Significant
CDK4_MDM2_CCND1
TP53_ATRX
SF3B1mut
AdjustedLog2OddsRatio
k
DFCI Discovery Cohort, n = 86
TP53/ATRX
SF3B1mut
MM Triple WT
CDK4/MDM2/CCND1
Adjusted Log2 Odds Ratio
l
GENIE Validation Cohort, n = 253

**Extended Data Fig. 3: a**, Number of patients contributed by each center within the AACR GENIE cohort. COLU = Columbia University; DFCI = Dana-Farber Cancer Institute; GRCC = Gustave Roussy Cancer Campus; JHU = Johns Hopkins University; MSK = Memorial Sloan Kettering Cancer Center; NKI = Netherlands Cancer Institute; PROV = Providence Health & Services; UCHI = University of Chicago; UCSF = University of California, San Francisco; UHN = University Health Network; VHIO = Vall d'Hebron Institute of Oncology; VLCC = Victorian Life Sciences Computation Initiative; WAKE = Wake Forest University; YALE = Yale University. **b**,**c**, Distribution of tumor sites and disease status in the DFCI (**b**) and AACR GENIE (**c**) cohorts. **d**, The overlap between top 10% altered genes in the DFCI and AACR GENIE cohorts with driver genes. **e**,**f**, Co-mutational plot of targeted sequencing data from the DFCI (**e**; $n$ = 86) and AACR GENIE (**f**; $n$ = 253) cohorts showing alterations in driver genes. Each column represents one patient, and each row represents one gene. Colors indicate alteration type. HA = high amplification; 2DEL = loss of both copies; Indel = insertions or deletions; Struc Var=structural variation. **g**,**h**, Heat map of pairwise gene associations for genes altered in > 5% of patients and gene pairs with OR > 3 or OR < 0.25 in DFCI (**g**) and AACR GENIE (**h**) cohorts. **i**, Permutation analysis of mutual exclusivity TP53 and ATRX ($P = 9.9 \times 10^{-4}$); and SF3B1 and KIT ($P = 9.9 \times 10^{-4}$) in AACR GENIE cohort ($n$ = 203). **j**, Co-occurrence between MM Triple WT group and alterations in other driver genes. **k**,**l**, Heatmaps of mutual exclusivity analysis among MM genetic subtypes in the DFCI (**k**) and AACR GENIE (**l**) cohorts. Heatmap is colored as per adjusted Log2(OR), and corresponding numerical values are depicted in each block.

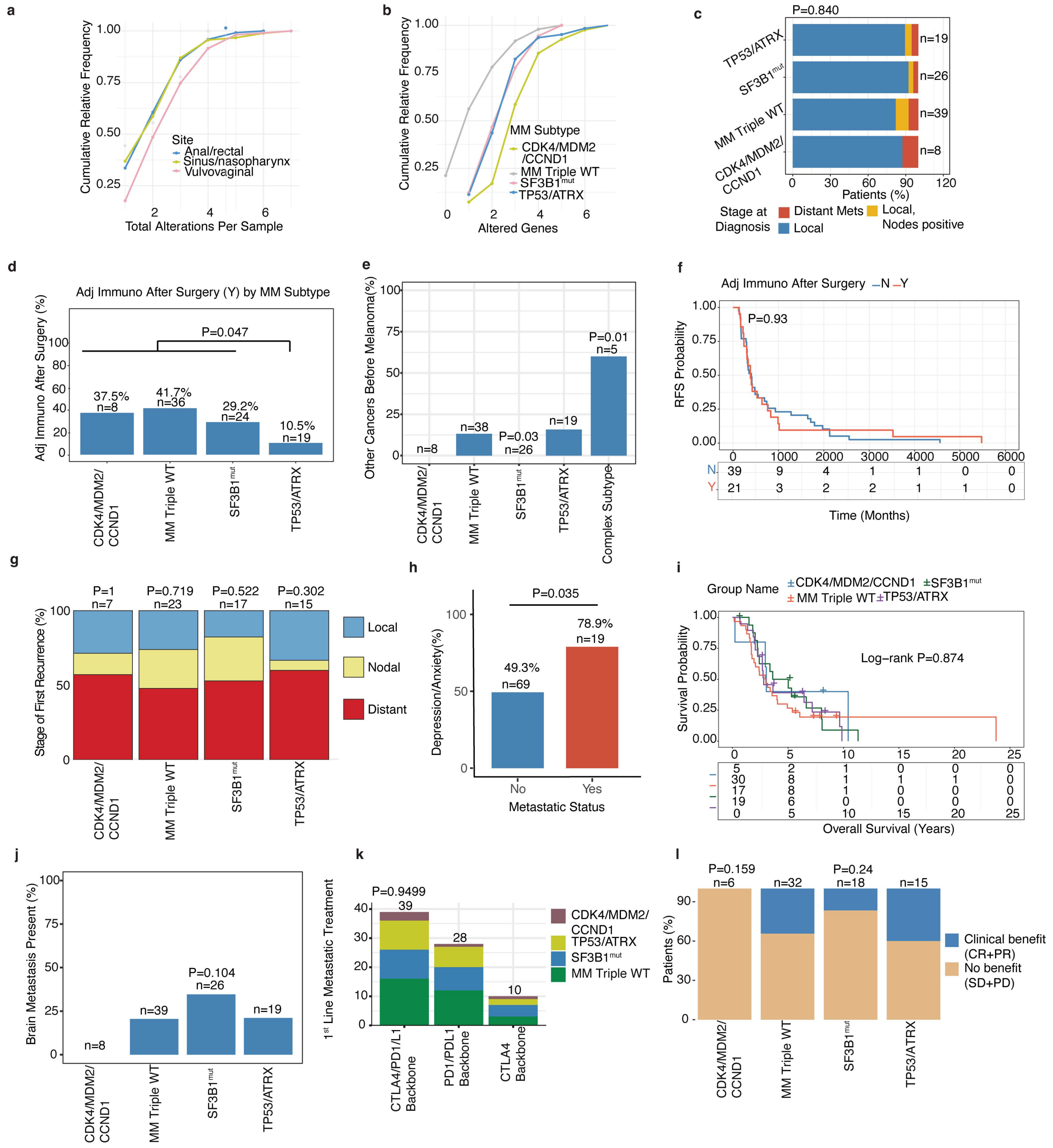

a
Cumulative Relative Frequency
Total Alterations Per Sample
Site
Anal/rectal
Sinus/nasopharynx
Vulvovaginal
b
Cumulative Relative Frequency
Altered Genes
MM Subtype
CDK4/MDM2 /CCND1
MM Triple WT
SF3B1mut
TP53/ATRX
c
P=0.840
TP53/ATRX
SF3B1mut
MM Triple WT
CDK4/MDM2/ CCND1
n=19
n=26
n=39
n=8
Patients (%)
Stage at Diagnosis
Distant Mets
Local, Nodes positive
Local
d
Adj Immuno After Surgery (Y) by MM Subtype
Adj Immuno After Surgery (%)
P=0.047
37.5% n=8
41.7% n=36
29.2% n=24
10.5% n=19
CDK4/MDM2/ CCND1
MM Triple WT
SF3B1mut
TP53/ATRX
e
Other Cancers Before Melanoma(%)
n=8
n=38
P=0.03 n=26
n=19
P=0.01 n=5
CDK4/MDM2/ CCND1
MM Triple WT
SF3B1mut
TP53/ATRX
Complex Subtype
f
Adj Immuno After Surgery —N —Y
RFS Probability
P=0.93
N 39 9 4 1 1 0 0
Y 21 3 2 2 1 1 0
Time (Months)
g
P=1 n=7
P=0.719 n=23
P=0.522 n=17
P=0.302 n=15
Stage of First Recurrence (%)
Local
Nodal
Distant
CDK4/MDM2/ CCND1
MM Triple WT
SF3B1mut
TP53/ATRX
h
P=0.035
Depression/Anxiety(%)
49.3% n=69
78.9% n=19
No
Yes
Metastatic Status
i
Group Name
CDK4/MDM2/CCND1
SF3B1mut
MM Triple WT
TP53/ATRX
Survival Probability
Log−rank P=0.874
5 2 1 0 0 0
30 8 1 1 1 0
17 8 1 0 0 0
19 6 0 0 0 0
Overall Survival (Years)
j
Brain Metastasis Present (%)
n=8
n=39
P=0.104 n=26
n=19
CDK4/MDM2/ CCND1
MM Triple WT
SF3B1mut
TP53/ATRX
k
P=0.9499
39
28
10
1st Line Metastatic Treatment
CDK4/MDM2/ CCND1
TP53/ATRX
SF3B1mut
MM Triple WT
CTLA4/PD1/L1 Backbone
PD1/PDL1 Backbone
CTLA4 Backbone
l
P=0.159 n=6
n=32
P=0.24 n=18
n=15
Patients (%)
Clinical benefit (CR+PR)
No benefit (SD+PD)
CDK4/MDM2/ CCND1
MM Triple WT
SF3B1mut
TP53/ATRX

**Extended Data Fig. 4: a**, Cumulative gene alteration frequency across tumor sites (Kolmogorov–Smirnov test). **b**, Cumulative gene alteration frequency across MM genetic subtypes (Kolmogorov–Smirnov test). **c**, Metastatic stage at diagnosis by MM subtype (Chi-square test). **d**, Adjuvant therapy received by each MM subtype with significant Fisher's exact *P* values. **e**, History of prior malignancies before melanoma diagnosis by MM genetic subtype, with significant Fisher's exact *P* values shown. **f**, Effect of adjuvant therapy on recurrence-free survival (RFS) using the overall log-rank test. **g**, Type of first recurrence according to MM genetic subtype (Chi-square test). **h**, Depression and anxiety according to metastatic status (Fisher's exact test). **i**, Overall survival by MM genetic subtype with overall log-rank test. **j**, Brain metastasis in MM genetic subtypes; a significant Fisher's exact *P* value is shown. **k**, First-line of metastatic therapy type received by each genetic subtype; colors indicate therapy types. **l**, Clinical response to the first-line of metastatic therapy in different genetic subtypes CR = complete response; PR = partial response; PD = progressive disease; SD = stable disease (Chi-square test). **m**, Clinical outcomes of first-line metastatic therapy in MM genetic subtypes (Chi-square test).

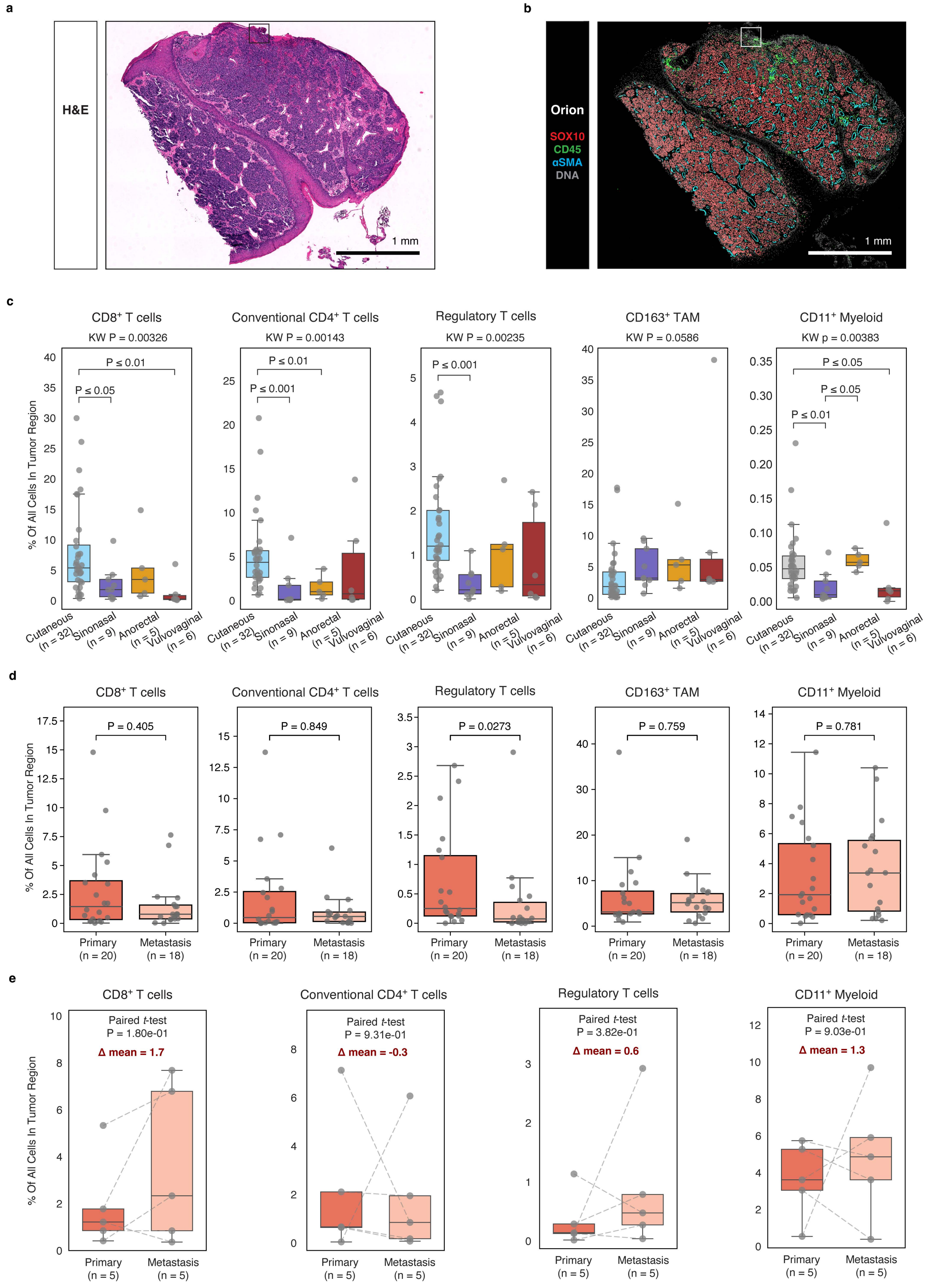
a
H&E
1 mm
b
Orion
SOX10
CD45
αSMA
DNA
1 mm
c
CD8+ T cells
KW P = 0.00326
P ≤ 0.01
P ≤ 0.05
Conventional CD4+ T cells
KW P = 0.00143
P ≤ 0.01
P ≤ 0.001
Regulatory T cells
KW P = 0.00235
P ≤ 0.001
CD163+ TAM
KW P = 0.0586
CD11+ Myeloid
KW p = 0.00383
P ≤ 0.05
P ≤ 0.05
P ≤ 0.01
% Of All Cells In Tumor Region
Cutaneous (n = 32)
Sinonasal (n = 9)
Anorectal (n = 5)
Vulvovaginal (n = 6)
d
CD8+ T cells
P = 0.405
Conventional CD4+ T cells
P = 0.849
Regulatory T cells
P = 0.0273
CD163+ TAM
P = 0.759
CD11+ Myeloid
P = 0.781
% Of All Cells In Tumor Region
Primary (n = 20)
Metastasis (n = 18)
e
CD8+ T cells
Paired t-test
P = 1.80e-01
Δ mean = 1.7
Conventional CD4+ T cells
Paired t-test
P = 9.31e-01
Δ mean = -0.3
Regulatory T cells
Paired t-test
P = 3.82e-01
Δ mean = 0.6
CD11+ Myeloid
Paired t-test
P = 9.03e-01
Δ mean = 1.3
% Of All Cells In Tumor Region
Primary (n = 5)
Metastasis (n = 5)

**Extended Data Fig. 5: a**, H&E-staining imaging of MM sample. The region outlined with a black rectangle corresponds to the magnified region shown in **Fig. 3b**. Scale bars, 1 mm. **b**, Orion multiplexed immunofluorescence imaging of the same MM sample, stained for SOX10 (red), CD45 (green), aSMA (cyan) and DNA (grey). The region outlined with a white rectangle corresponds to the magnified region shown in **Fig. 3b**. Scale bars, 1 mm. **c**, Quantification of tumor-infiltrating immune cell populations in primary CM ($n$ = 32) and in primary MM ($n$ = 20) by anatomic subsite: sinonasal ($n$ = 9), anorectal ($n$ = 5) and vulvovaginal ($n$ = 6). Shown are the proportions of $CD8^+$ T cells, conventional $CD4^+$ T cells, regulatory T cells, $CD163^+$ TAMs, and $CD11c^+$ myeloid cells within the tumor compartment. Box plots show the median and interquartile range (IQR), with whiskers extending to 1.5 × IQR and individual tumors shown as points. Groups were compared using two-sided Mann-Whitney $U$ tests; significant $P$ values ($P < 0.05$) are shown. **d**, Quantification of tumor-infiltrating immune cell populations in unpaired treatment naïve primary ($n$ = 20) and metastatic ($n$ = 18) MM. Shown are the proportions of $CD8^+$ T cells, conventional $CD4^+$ T cells, regulatory T cells, $CD163^+$ TAMs, and $CD11c^+$ myeloid cells within the tumor compartment. Box plots and statistics as in **c**. **e**, Quantification of tumor-infiltrating immune cell populations in paired treatment naïve primary and metastatic MM ($n$ = 5 pairs). Shown are the proportions of $CD8^+$ T cells, conventional $CD4^+$ T cells, regulatory T cells, and $CD11c^+$ myeloid cells within the tumor compartment. Dashed lines connect paired samples. Groups were compared using the parried student $t$-test.

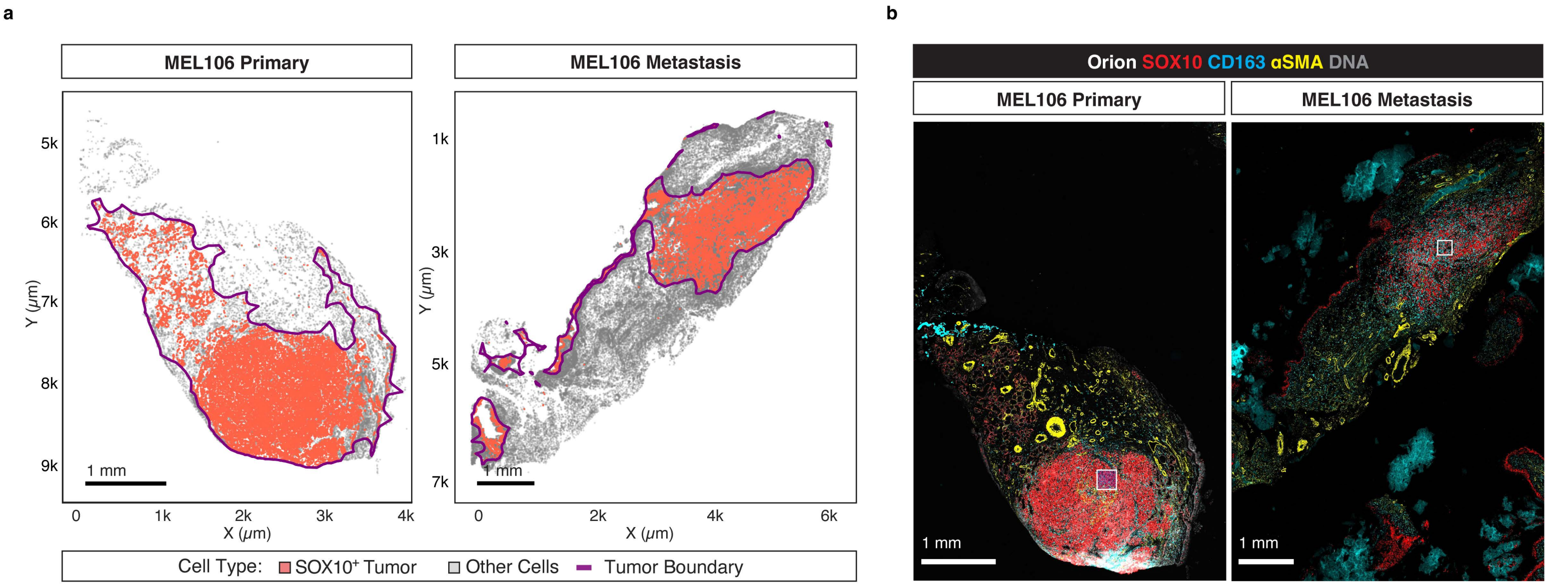
a
MEL106 Primary
MEL106 Metastasis
5k
6k
7k
8k
9k
Y (μm)
1 mm
0
1k
2k
3k
4k
X (μm)
1k
3k
5k
7k
6k
Cell Type: SOX10+ Tumor
Other Cells
Tumor Boundary
b
Orion SOX10 CD163 αSMA DNA
1 mm

**Extended Data Fig. 6: a**, Scatter plot showing the single-cell spatial maps of matched primary and metastatic MM tumors from the same patient (MEL106), derived from Orion multiplexed immunofluorescence images. $SOX10^{+}$ tumor cells are shown in red and all remaining cells in grey. Tumor boundaries (purple) were defined by spatial clustering of $SOX10^{+}$ tumor cells (Methods). Axes indicate spatial coordinates micrometers. Scale bars, 1 mm. **b**, Orion multiplexed immunofluorescence images of the MEL106 paired treatment-naive primary and metastatic MM tumors. Orion images are stained for SOX10 (red), CD163 (cyan), aSMA (yellow) and DNA (grey). The region outlined by a white rectangle corresponds to the magnified region shown in **Fig. 3d**. Scale bars, 1 mm.

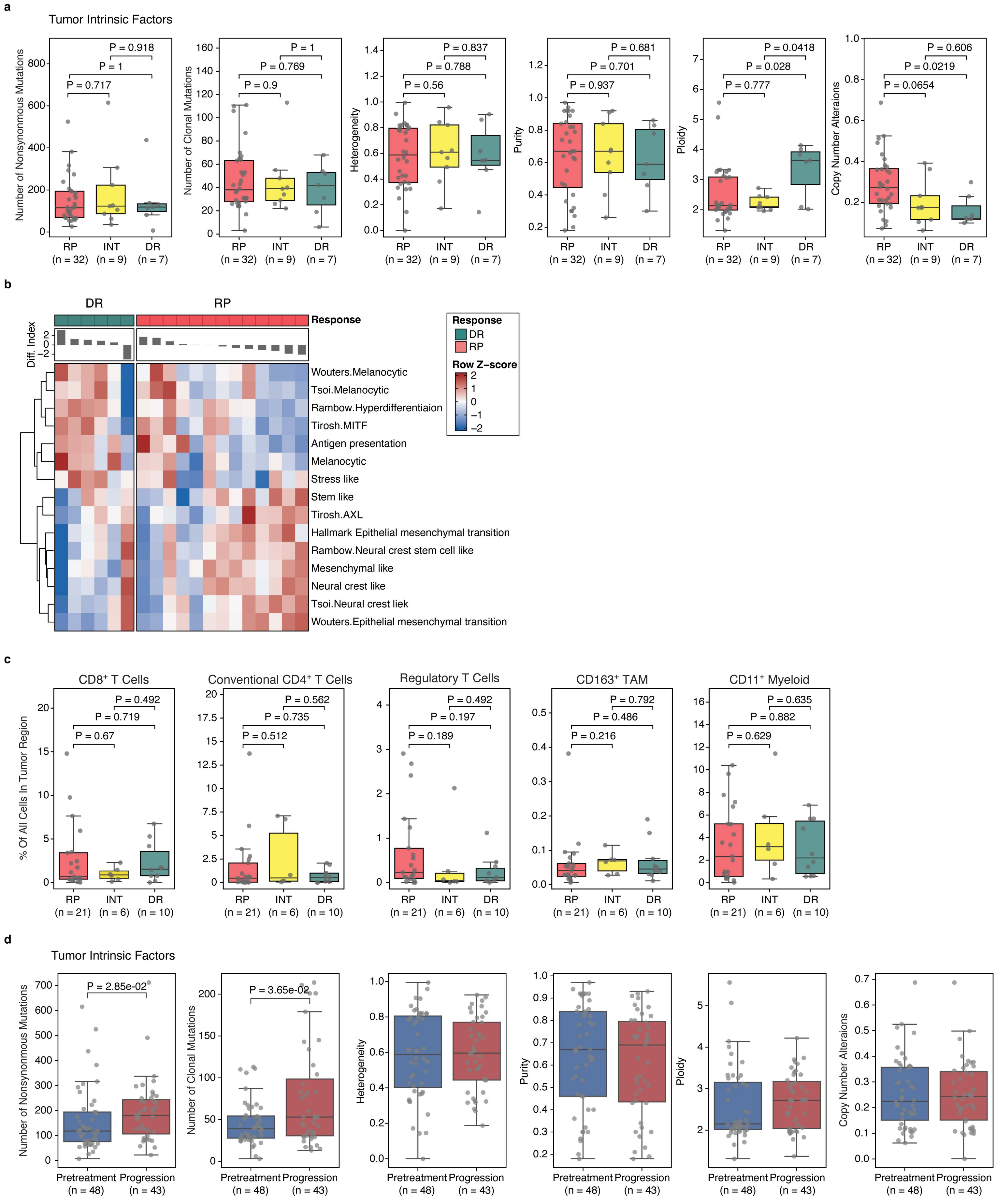
a
Tumor Intrinsic Factors
Number of Nonsynonymous Mutations
P = 0.918
P = 1
P = 0.717
Number of Clonal Mutations
P = 1
P = 0.769
P = 0.9
Heterogeneity
P = 0.837
P = 0.788
P = 0.56
Purity
P = 0.681
P = 0.701
P = 0.937
Ploidy
P = 0.0418
P = 0.028
P = 0.777
Copy Number Alteraions
P = 0.606
P = 0.0219
P = 0.0654
RP (n = 32)
INT (n = 9)
DR (n = 7)
b
DR
RP
Response
Diff. Index
Wouters.Melanocytic
Tsoi.Melanocytic
Rambow.Hyperdifferentiaion
Tirosh.MITF
Antigen presentation
Melanocytic
Stress like
Stem like
Tirosh.AXL
Hallmark Epithelial mesenchymal transition
Rambow.Neural crest stem cell like
Mesenchymal like
Neural crest like
Tsoi.Neural crest liek
Wouters.Epithelial mesenchymal transition
Response
DR
RP
Row Z-score
c
CD8⁺ T Cells
% Of All Cells In Tumor Region
P = 0.492
P = 0.719
P = 0.67
Conventional CD4⁺ T Cells
P = 0.562
P = 0.735
P = 0.512
Regulatory T Cells
P = 0.492
P = 0.197
P = 0.189
CD163⁺ TAM
P = 0.792
P = 0.486
P = 0.216
CD11⁺ Myeloid
P = 0.635
P = 0.882
P = 0.629
RP (n = 21)
INT (n = 6)
DR (n = 10)
d
Tumor Intrinsic Factors
Number of Nonsynonymous Mutations
P = 2.85e-02
Number of Clonal Mutations
P = 3.65e-02
Heterogeneity
Purity
Ploidy
Copy Number Alteraions
Pretreatment (n = 48)
Progression (n = 43)

**Extended Data Fig. 7: a**, Genomic features comparison among DR (*n* = 7), INT (*n* = 9), and RP (*n* = 32) pretreatment MM tumors. Shown are number of nonsynonymous mutations, number of clonal mutations, heterogeneity, purity, ploidy, and the proportion of copy-number alterations derived from WES. Box plots show the median and interquartile range (IQR), with whiskers extending to 1.5 × IQR and individual tumors shown as points. Groups were compared using two-sided Mann-Whitney U tests; significant *P* values ($P < 0.05$) are shown. **b**, Heatmap of melanoma tumor states signature scores in pretreatment DR (*n* = 6) and RP (*n* = 13) tumors. Signature scores are row-wise z-scores; rows are clustered by Euclidean distance with optimal leaf ordering. Samples are ordered within each response group by a differentiation index (top bar), defined as the mean z-score of melanocytic signatures minus the mean z-score of dedifferentiated signatures. **c**, Quantification of tumor-infiltrating immune cell populations in pretreatment DR (*n* = 10), INT (*n* = 6) and RP (*n* = 21) tumors. Shown are the proportions of $CD8^+$ T cells, conventional $CD4^+$ T cells, regulatory T cells, $CD163^+$ TAMs and $CD11c^+$ myeloid cells within the tumor compartment. Box plots show the median and IQR, with whiskers extending to 1.5 × IQR and individual tumors shown as points. Groups were compared using two-sided Mann-Whitney *U* tests. **d**, Genomic features comparison between pretreatment (*n* = 48) and progression (*n* = 43) MM tumors. Shown are nonsynonymous mutation burden, clonal mutation burden, intratumoral heterogeneity, tumor purity, ploidy and aneuploidy. Box plots show the median and IQR, with whiskers extending to 1.5 × IQR and individual tumors shown as points. Groups were compared using two-sided Mann-Whitney *U* tests; *P* values are shown in the figure.

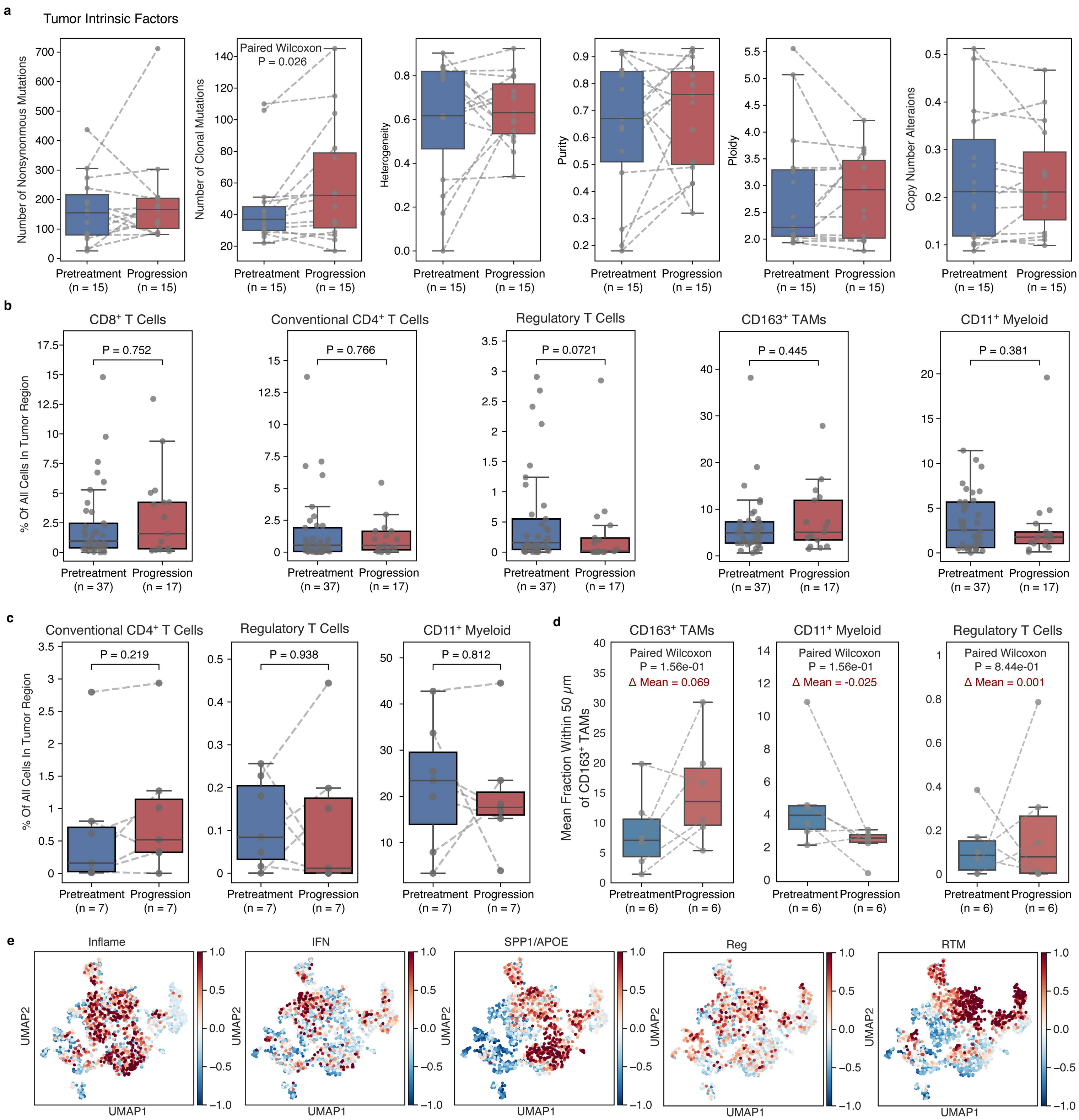

a
Tumor Intrinsic Factors
Number of Nonsynonymous Mutations
Paired Wilcoxon
P = 0.026
Number of Clonal Mutations
Heterogeneity
Purity
Ploidy
Copy Number Alterations
Pretreatment (n = 15)
Progression (n = 15)
b
CD8+ T Cells
P = 0.752
Conventional CD4+ T Cells
P = 0.766
Regulatory T Cells
P = 0.0721
CD163+ TAMs
P = 0.445
CD11+ Myeloid
P = 0.381
% Of All Cells In Tumor Region
Pretreatment (n = 37)
Progression (n = 17)
c
Conventional CD4+ T Cells
P = 0.219
Regulatory T Cells
P = 0.938
CD11+ Myeloid
P = 0.812
% Of All Cells In Tumor Region
Pretreatment (n = 7)
Progression (n = 7)
d
CD163+ TAMs
Paired Wilcoxon
P = 1.56e-01
Δ Mean = 0.069
CD11+ Myeloid
Paired Wilcoxon
P = 1.56e-01
Δ Mean = -0.025
Regulatory T Cells
Paired Wilcoxon
P = 8.44e-01
Δ Mean = 0.001
Mean Fraction Within 50 μm of CD163+ TAMs
Pretreatment (n = 6)
Progression (n = 6)
e
Inflame
IFN
SPP1/APOE
Reg
RTM
UMAP1
UMAP2

**Extended Data Fig. 8: a,** Box plots showing paired comparison of genomic features between pretreatment and post-progression MM tumors from the same patient (*n* = 15 pairs). Boxes show the median and interquartile range (IQR), with whiskers extending to 1.5 × IQR; dashed lines connect samples from the same patient. Groups were compared using two-sided Wilcoxon signed-rank tests; only significant comparisons are annotated. **b**, Quantification of tumor-infiltrating immune cell populations in pretreatment (*n* = 37) and post-progression (*n* = 17) tumors (unpaired analysis). Shown are the proportions of $CD8^+$ T cells, conventional $CD4^+$ T cells, regulatory T cells, $CD163^+$ TAMs and $CD11c^+$ myeloid cells within the tumor compartment. Box plots show the median and IQR, with whiskers extending to 1.5 × IQR and individual tumors shown as points. Groups were compared using two-sided Mann-Whitney *U* tests. **c**, Paired comparison of tumor-infiltrating immune cell populations between pretreatment and post-progression MM tumors from the same patient (*n* = 7 pairs). Dashed lines connect samples from the same patient. Groups were compared using two-sided Wilcoxon signed-rank tests. **d**, Spatial neighborhood analysis of $CD163^+$ TAMs in paired pretreatment and post-progression MM tumors (*n* = 6 pairs). For each sample, $CD163^+$ TAMs were used as the central cell population and the mean fraction of the indicated cell type within a 50-μm radius was quantified. Dashed lines connect samples from the same patient; Δmean indicates the mean paired difference (progression − pretreatment). Groups were compared using two-sided Wilcoxon signed-rank tests. **e,** UMAP embeddings of macrophages colored by signature scores for the indicated tumor-associated macrophage programs (Inflame, IFN, SPP1/APOE, Reg, RTM). Color indicates the scaled module score, capped at ± 1.
